# The influence of study characteristics on coordinate-based fMRI meta-analyses

**DOI:** 10.1101/144071

**Authors:** Han Bossier, Ruth Seurinck, Simone Kühn, Tobias Banaschewski, Gareth J. Barker, Arun L.W. Bokde, Jean-Luc Martinot, Herve Lemaitre, Tomáš Paus, Sabina Millenet, Beatrijs Moerkerke, The IMAGEN Consortium

## Abstract

Given the increasing amount of neuroimaging studies, there is a growing need to summarize published results. Coordinate-based meta-analyses use the locations of statistically significant local maxima with possibly the associated effect sizes to aggregate studies. In this paper, we investigate the influence of key characteristics of a coordinate-based meta-analysis on (1) the balance between false and true positives and (2) the reliability of the outcome from a coordinate-based meta-analysis. More particularly, we consider the influence of the chosen group level model at the study level (fixed effects, ordinary least squares or mixed effects models), the type of coordinate-based meta-analysis (Activation Likelihood Estimation, fixed effects and random effects meta-analysis) and the amount of studies included in the analysis (10, 20 or 35). To do this, we apply a resampling scheme on a large dataset (*N* = 1400) to create a test condition and compare this with an independent evaluation condition. The test condition corresponds to subsampling participants into studies and combine these using meta-analyses. The evaluation condition corresponds to a high-powered group analysis. We observe the best performance when using mixed effects models in individual studies combined with a random effects meta-analysis. This effect increases with the number of studies included in the meta-analysis. We also show that the popular Activation Likelihood Estimation procedure is a valid alternative, though the results depend on the chosen threshold for significance. Furthermore, this method requires at least 20 to 35 studies. Finally, we discuss the differences, interpretations and limitations of our results.

## 1 Introduction

Over the past two decades, there has been a substantial increase in the number of functional Magnetic Resonance Imaging (fMRI) studies, going from 20 publications in 1994 to over 5000 in 2015. Despite this vast amount of fMRI literature, it remains challenging to establish scientific truth across these often-contradictory studies.

First, fMRI studies tend to have small sample sizes to detect realistic effect sizes (median estimated sample size in 2015 = 28.5; Poldrack et al., 2017) as among other causes scanning participants is costly and time consuming. The large multiple testing problem and ensuing corrections make statistical testing in fMRI conservative, thereby further reducing statistical power or probability to detect true activation (Durnez et al., 2014; Lieberman and Cunningham, 2009). As a consequence, the probability that a statistically significant effect reflects true activation is reduced (Button et al., 2013). This can lead to more false negatives (missing true activation) as well as more false positives (detecting activation where there is none) in published fMRI studies. Second, the diversity of pre-processing steps and analysis pipelines have made fMRI studies challenging to replicate (Carp, 2012b, a), even though researchers recognize the value of both reproducibility (obtaining identical parameter estimates compared to the original experiment using the same analysis and data; Poldrack and Poline, 2015) and replicability (the ability of an entire experiment to be replicated by gathering new data using the exact same materials and methods; Patil et al., 2016). Roels et al. (2015) also showed there is variability in the number of significant features depending on the data-analytical methods used.

Several approaches have been offered to overcome these challenges. A first remediating step is to promote transparency, pre-registration and open science initiatives such as data sharing or using standardized protocols in organizing and managing data (Pernet and Poline, 2015; Gorgolewski et al., 2016; Gorgolewski and Poldrack, 2016; Poline et al., 2012; Poldrack et al., 2017). A second approach to establish scientific truth across studies, is to accumulate knowledge by scientifically combining previous results using meta-analysis (Lieberman and Cunningham, 2009; Yarkoni et al., 2010). Combining findings across studies increases power to detect true effects, while false positives are not expected to replicate across studies, given a representative set of unbiased results. Furthermore, meta-analyses can generate new scientific questions (Wager et al., 2009).

Originally, meta-analyses were developed to aggregate single univariate effect sizes (Borenstein et al., 2009). In an individual fMRI study however, the brain is divided in a large amount of artificially created cubes (voxels). Until recently, the standard approach was to only report coordinates in 3D space of peaks of activity that survive a statistical threshold. These are called foci, peaks or local maxima. While guidelines are shifting towards making statistical maps or full data sets of a study available, many findings in the literature only consist of locations of activation. In these cases, an fMRI meta-analysis is limited to those voxels for which information is at hand. This is termed a coordinate-based meta-analysis (CBMA, see e.g. Paus 1996; Paus et al. 1998). When full images (and hence information in all voxels) are available, methods designed for image-based meta-analysis (IBMA) can be used (Radua and Mataix-Cols, 2012; Salimi-Khorshidi et al., 2009).

In this study, we focus on CBMA for which different algorithms exist (Wager et al., 2007; Radua and Mataix-Cols, 2012). In particular, we consider the popular Activation Likelihood Estimation (ALE) (Turkeltaub et al., 2002, 2012) and effect size based methods such as seed based d-mapping (SBdM, formerly called effect size-signed differential mapping) (Radua et al., 2012).

The ALE algorithm considers a reported local maximum as a center of a spatial probability distribution. As such, the method only requires the location of the peak and then searches for brain regions where spatial convergence can be distinguished from random clustering of peaks.

Effect size based methods on the other hand transform t-values of reported local maxima are transformed into effect size estimates and calculate a weighted average of the reported evidence. The weights determine the underlying meta-analysis model. For instance, the weights in seed based *d*-mapping include within-study and between-study variability which corresponds to a random effects model. If the weights ignore the between-study variability one obtains a fixed effects model.

In this paper, we evaluate the influence of study characteristics on the statistical properties of CBMA techniques for fMRI. Previous work by Eickhoff et al. (2016b) and Radua et al. (2012) already evaluated statistical properties of CBMA algorithms or tested software for implementation errors (Eickhoff et al., 2016a). However, these studies did not study the effect of input characteristics at the individual study level on the performance of these CBMA algorithms.

We investigate the influence of the group level model on the performance of various CBMA procedures. More specifically, we test the effect of pooling subjects at the individual study level using either a fixed effects, ordinary least squares (OLS) or mixed effects group level model on the outcome of the meta-analyses methods mentioned above. As in Eickhoff et al. (2016b) we also evaluate the effect of the number of studies in the meta-analysis (*K*). Extending on their work, we consider the case for *K* = 10, 20 and 35 when using ALE as well as effect size based CBMA’s using a fixed and random effects model. We consider two performance measures: the balance between false positives and true positives and the activation reliability as a proxy for replicability.

We approach this problem by applying a resampling scheme on a large dataset from the IMAGEN project (Schumann et al., 2010) and create meta-analyses (i.e. test conditions) which we compare against a high powered large sample size study as a reference (i.e. an evaluation condition).

In the following section, we discuss the dataset, give a theoretical overview of the three models to pool subjects at study level and discuss the three models for coordinate-based meta-analysis. In the next sections, we present the design of the study with the chosen performance measures and discuss our findings.

## 2 Materials and Methods

The code containing the design and analysis of the results in this paper are available at: https://github.com/NeuroStat/PaperStudyCharCBMA

### 2.1 Data

We use preprocessed data from the IMAGEN project (Schumann et al., 2010). This is a large genetic-neuroimaging study on reinforcement-related behaviour in adolescents with the goal to identify its predictive value for the development of frequent psychiatric disorders across Europe. The database contains fMRI data from 1487 adolescents aged between 13 and 15 years, acquired across several research centers on 3 Tesla scanners from different manufactures. The data are stored and preprocessed at the Neurospin center using SPM8 (http://www.fil.ion.ucl.ac.uk/spm/software/spm8/).

The scanning sessions of interest involved a global cognitive assessment. Note that we only use a part of the entire IMAGEN database for which data was acquired using the following tasks. In a fast-event related design, participants had to do a series of alternating cognitive/motor tasks. Two of these are (1) reading sentences in silence and (2) solving math subtractions in silence. These math questions were single digits (0-9) that had to be subtracted from a digit between 11-20. Each of these trials was presented for 10 times with a probabilistic inter-stimulus interval of on average 3 seconds (see also Pinel et al., 2007). We use the contrast MATH > LANGUAGE (2 - 1) for this study.

A BOLD time series was recorded for each participant using echoplanar imaging with an isotropic voxel size of 3.4 mm, isotropic and temporal resolutions of 2.2 seconds. A total of 160 volumes are obtained. For each participant, a structural T1-weighted image (based on the ADNI protocols (http://adni.loni.usc.edu/methods/documents/mri-protocols/)) was acquired for registration.

Preprocessing included slice-timing correction, movement correction, coregistration to the segmented structural T1-weighted images, non-linear warping on the MNI space using a custom EPI template and spatial smoothing of the signal with a 5 mm Gaussian Kernel (Imagen fMRI data analysis methods, revision2, July 2010).

In the first level analysis, all experimental manipulations were modelled using a general linear model with a standard autoregressive (AR(1)) noise model and 18 estimated movement parameters as nuisance terms. This resulted in a statistical map for each parameter estimate and a map reflecting the residual variance of the model fit.

In this study, we use for each participant (1) the contrast map or the difference between the parameter estimate maps for MATH and LANGUAGE and (2) an error map for that contrast derived from the residual variance map. After visual inspection for errors or artefacts we removed 87 participants from which parts of the brain were missing. To automate, we used a cut-off corresponding to 96% of the median number of masked voxels over all subjects in the database.

### 2.2 Group level models

Localizing significant brain activity in an fMRI data-analysis is based on the statistical parametric map of contrasting conditions associated with all participants involved in an experiment. In this study, we focus on the univariate approach in which activation is tested in a voxelwise manner through general linear models (GLMs). Due to computational constraints, the analysis is typically executed in a two stage GLM procedure (Beckmann et al., 2003). In a first step, the measured time series (BOLD signal) of each subject is modelled by a linear combination of nuisance terms and the expected time series under the experimental design. Note that such a model is fitted for each voxel *ν* (*ν* = 1,…, *S*) separately. In what follows, we drop the index *ν* for ease of notation. This first stage model for a single subject *i* (*i* = 1,…,*N*) can be written as follows (Friston et al., 1995):

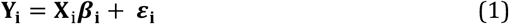

where **Y_i_** is a one dimensional vector of length *T* containing the measurements of the BOLD signal on *T* different time points, **X**_i_ is a matrix of dimension *T* × *p* that contains a convolution of the stimulus onset function with a hemodynamic response function (HRF; see e.g. Henson and Friston, 2007) as well as possible nuisance covariates, ***β*_i_** = (*β*_*i*1_,…,*β_ip_*) is a vector of parameter estimates and ***ε**_i_* is a one dimensional vector of length *T* containing the within-subject random error with mean zero. Temporal correlation is removed through prewhitening. Localizing activation proceeds by testing specific contrasts of ***β_i_***. Let *c* represent a contrast vector, the null hypothesis *H*_0_ can then be expressed as: ***cβ_i_*** = 0. For inference, one typically assumes independent error terms that follow a Gaussian distribution.

In a second step, parameter estimates obtained at the first stage are combined over *N* subjects to obtain group level estimates. More particularly, we use the vector of estimated first level contrasts 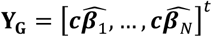. Then, for every voxel *ν* (*ν* = 1,…,*S*), we estimate the following model:

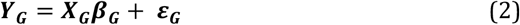

in which ***X*_G_** is a group design matrix and ***ε*_G_** a mixed-effects zero mean error component containing between subject variability and within subject variability. In the simplest case, we are interested in the average group activation. Therefore, when testing the null hypothesis *H*_0_ of no group activation (***β*_G_** = 0), **X_G_** is a column vector of length *N* with all elements equal to 1 and the test statistic is identical to a one-sample *t*-test:

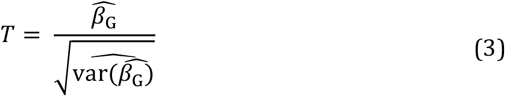

Under the assumption that 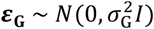 this test statistic follows a t-distribution under *H*_0_. Alternatively, it is possible to test differences between groups of subjects (e.g. patients versus controls) by incorporating additional regressors in the group design matrix. As statistical tests are performed in all voxels simultaneously, adjustments for multiple testing need to be imposed.

Several methods are available to estimate ***β*_G_** and var(***β*_G_**) in model (2). We consider the Ordinary Least Squares (OLS), Fixed Effects (FE) and Mixed Effects (ME) approaches. In this study, we use the FSL software library (Smith et al., 2004) and therefore only outline the implementation of these methods as described in Woolrich et al. (2004). For a discussion of different implementations in other software packages, see Mumford and Nichols (2006).

#### Ordinary Least Squares

In the OLS procedure (Holmes and Friston, 1998), one assumes that within subject variability is equal across all subjects (resulting in homogeneous residual variance). In the simple case of seeking group average activation, and as shown in Mumford and Nichols (2009), ***β*_G_** in model (2) can be estimated as 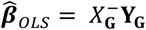 where - denotes the pseudo inverse. The residual error variance 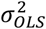 is estimated as 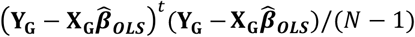, and therefore var(***β_OLS_***) can be estimated as 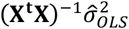. Under the assumption of Gaussian distributed error terms, the resulting test is equal to a one-sample t-test with *N* – 1 degrees of freedom (dof) on the contrast of parameter estimates **Y_G_** obtained at the first level.

In FSL, this model is termed *mixed effects: simple OLS*.

#### Fixed and mixed effects

Both for the fixed and mixed effects models, β_G_ in model (2) and var(β_G_) are estimated as follows:

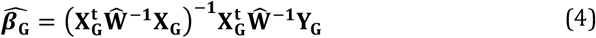

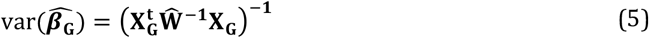

with **W** a weighting matrix. As is the case for OLS, the error terms in model 2 are typically assumed to follow a Gaussian distribution.

In the fixed effects model, the weights in **W** correspond to the within subject variability only (ignoring between subject variability). Hence, **W** is an N × N matrix equal to:

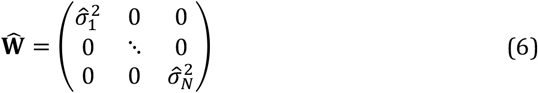

Thus, ***β*_G_** is equal to a weighted average of the first level contrast parameters with the weights corresponding to the inverse of the within subject variances.

These variances are easily estimated at the first level of the GLM procedure. The number of degrees of freedom in the fixed effects model depends on the number of scans per subject and the sample size at the second level (though FSL restricts the number of dof to a maximum of 1000 and is set equal to 999 when no information on the number of scans at the first level is provided).

In FSL, this model is termed *fixed effects*.

For the mixed effects model, between subject variability 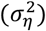 is incorporated into the weighting matrix:

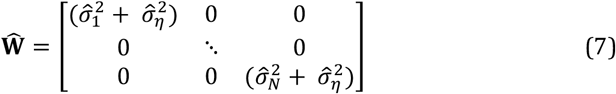

Estimating the variance components of the mixed effects model is complicated as (1) multiple components need to be estimated and (2) there are typically only a few measurements on the second level to estimate 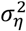. FSL relies on a fully Bayesian framework with reference priors (Woolrich et al., 2004). Inference on ***β*_G_** in model (2) then depends on its posterior distribution, conditional on the observed data (Mumford and Nichols, 2006). As suggested in Woolrich et al. (2004), a fast approximation is used first and then on voxels close to significance thresholding a slower Markov-Chain-Monte-Carlo sampling framework is applied to estimate all parameters of interest. The posterior marginal distribution of ***β*_G_** is assumed to approximate a multivariate t-distribution with noncentrality parameter 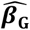. A lower bound on the number of degrees of freedom (i.e. *N* − *p_G_* with *p*_G_ the amount of parameters in the group design matrix **X_G_**) is used for the voxels with a test statistic close to zero and an EM algorithm (Dempster et al., 1977) is employed to estimate the effective degrees of freedom in voxels that are close to the significance threshold.

In FSL, this model is termed *mixed effects: FLAME1+2*.

### 2.3 Coordinate-based meta-analyses

#### 2.3.1 ALE

Coordinate based meta-analyses combine coordinates from several studies to assess convergence of the location of brain activation. The ALE algorithm (Turkeltaub et al., 2002, 2012) starts by creating an activation probability map for each study in the meta-analysis. The location of each reported peak in a study is modelled using a Gaussian kernel to reflect the spatial uncertainty of the peak activation. Voxels where kernels overlap due to multiple nearby peaks take the maximum probability. Next an ALE map is calculated by taking the voxelwise union of the probabilities over all studies. If *p_vm_* is the probability of a peak at voxel *ν* (*ν* = 1,…, *S*) in a study *m* (*m* = 1,…, *K*), then the union is defined as: 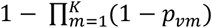.

A null distribution created with non-linear histogram integration is used for uncorrected voxel-level inference under the assumption of spatial independence (Eickhoff et al., 2012). Various corrections for multiple comparisons are available in ALE, but based on the large-scale simulation study in Eickhoff et al. (2016b), cluster-level family-wise error (cFWE) correction is preferred as it provides the highest power to detect a true underlying effect while being less susceptible to spurious activation in the meta-analysis. All ALE calculations were implemented using MATLAB scripts which corresponds to the ALE algorithm as described in Eickhoff et al. (2009, 2012, 2016b) and Turkeltaub et al. (2012) provided to us by Prof. dr. Simon Eickhoff (personal communication).

#### 2.3.2 Random effects CBMA

An alternative approach is to use the associated t-values of reported peaks to estimate corresponding effect sizes, enabling a weighted average of these effect sizes. Depending on the weights, this results in a random or fixed effects meta-analysis model. To evaluate the performance of these effect size based methods, we use the seed based *d*-mapping algorithm (SBdM), as described in Radua et al. (2012). However, we have carefully replicated this algorithm in R (R Core Team, 2015) to efficiently develop a fixed effects meta-analysis implementation (see below). As we cannot exclude slightly divergent results compared to the standalone version of SBdM (http://www.sdmproject.com), we choose to refer to this implementation as random effects CBMA. We follow the guidelines for significance testing as described in Radua et al. (2012).

Unlike ALE, the method assigns effect sizes to voxels. These correspond to the standardized mean (for a one sample design) known as Hedges’ *g* (Hedges, 1981) obtained from the peak height *t_vm_* in study *m* (*m* = 1,…, *K*)and voxel *ν* (*ν* = 1,…, *S*). For a given peak with height *t_νm_* stemming from a one-sample *t*-test and *N_m_* subjects, the effect size *g_vνm_* and a correction factor *J_m_* is given by:

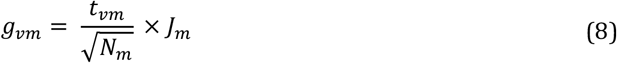

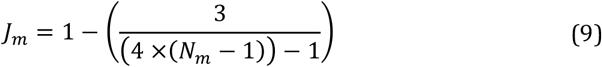

First, all coordinates of local maxima are smoothed using an unnormalized Gaussian kernel. The resulting map represents for each voxel the distance to a nearby peak. Effect sizes in voxels surrounding a peak are then obtained through multiplication of the peak effect size calculated using equation 8 and the smoothed map. The effect size in voxels where kernels overlap is an average weighted by the square of the distance to each nearby peak.

Once an effect size 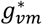 (i.e. the smoothed standardized effect size) is obtained in each voxel (which will be zero for voxels that are not near a peak), the variance of this effect size is obtained as follows (Hedges and Olkin, 1985):

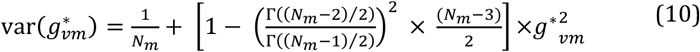

Combining all studies proceeds by calculating the weighted average *θ* through a random effects model:

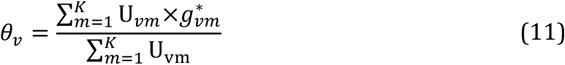

with the weights in U_*νm*_ being the inverse of the sum of both the within study variability (estimated using equation 10) and the between study variability (*τ*^2^). The latter is estimated through the DerSimonian & Laird estimator (DerSimonian and Laird, 1986).

In a final step, the null hypothesis H_0_: θ_v_ = 0 is calculated with the following Z-test: 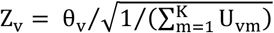 (Borenstein et al., 2009). A permutation approach with 20 iterations is used to create a combined null-distribution, in which each iteration is a whole brain permutation with close to 100,000 values. To optimally balance sensitivity and specificity, a threshold of *P* = 0.005 and *Z* > 1 is recommended, instead of classical multiple comparisons corrections (Radua et al., 2012). Since the effect size is imputed as 0 in voxels far from any peak, *Z* > 1 is a lot more unlikely under the empirical null distribution.

#### 2.3.3 Fixed effects CBMA

Finally, we also evaluate the performance of a fixed effects CBMA. This procedure only differs from the random effects CBMA with respect to the weights. A fixed effects model ignores heterogeneity across studies and only uses the within study variability to calculate the weights, U_vm_.

An illustration of ALE and an effect size based CBMA prior to thresholding can be seen in figure 1.

**Figure 1.**
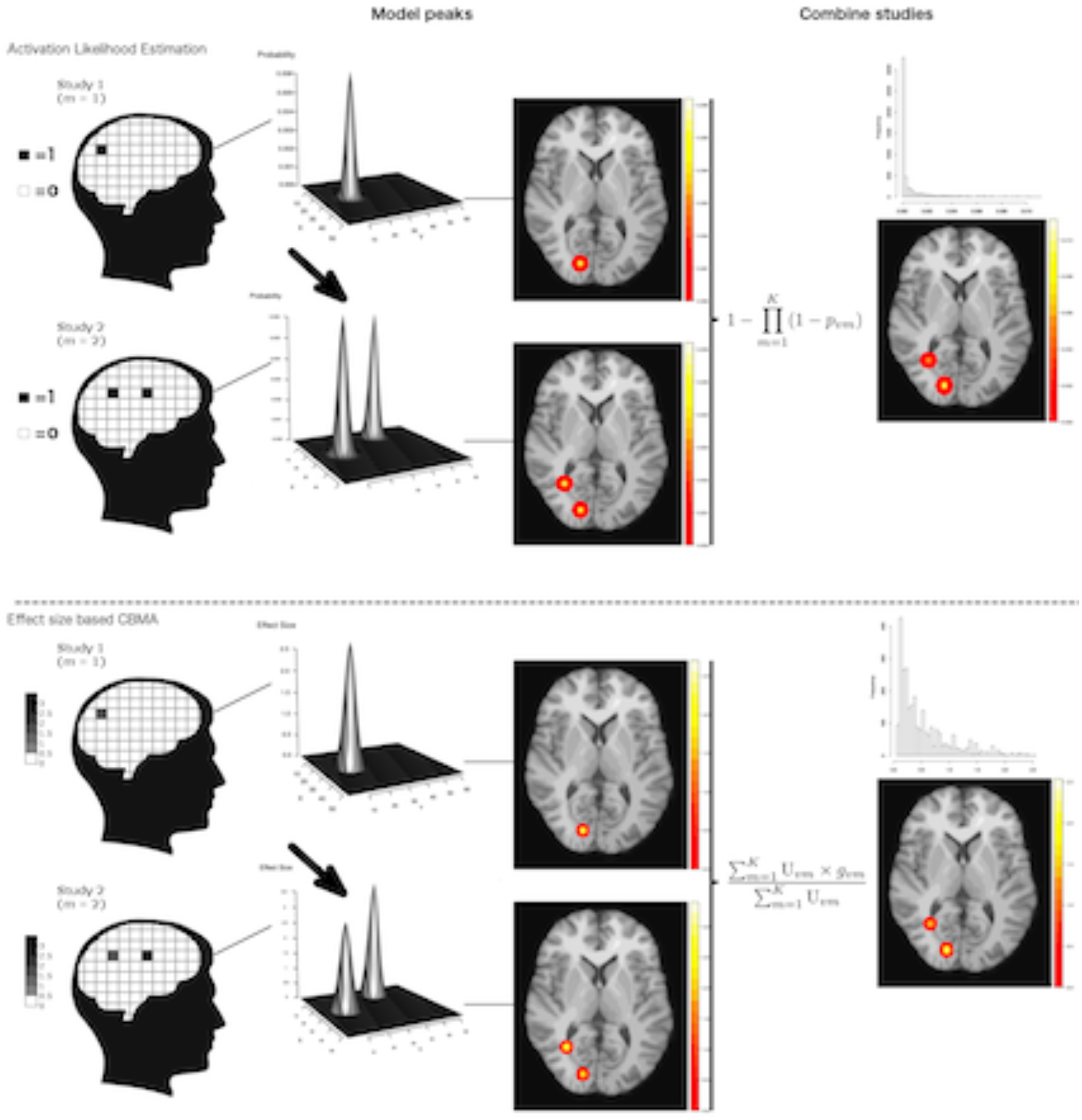
Illustration of ALE and an effect size based CBMA. Reported coordinates are first modelled by applying a Gaussian kernel. These are then combined either through calculating probabilities or by transforming the test-statistics to effect sizes and calculate a weighted average. Note that for illustration purpose, we only plot the values > 0 in the histograms. Illustration is prior to thresholding.

### 2.4 Design

In this section, we describe the set-up of our study to test the effect of pooling subjects at the individual study level on the outcome of methods for CBMA.

#### 2.4.1 Resampling scheme

The general study design is depicted in figure 2. In one iteration *l* (*l* = 1,…, *I*) or fold, *N_l_* subjects are sampled without replacement into an evaluation condition while *N_l_* different subjects go into a test condition. Next, the subjects in the test condition are subsampled into *K* smaller studies with varying sample sizes (mean = 20, SD = 5). No subsampling restriction into the *K* studies is imposed. However, to ensure independent results across iterations, we impose the restriction that subjects can only be used once in a test condition in one iteration and once in an evaluation condition in another iteration. Since this results in a trade-off between the number of iterations (*I*) and the number of subjects per iteration (*N_l_*), we consider three scenarios of the resampling scheme, allowing us to vary the number of studies in the meta-analyses in the test condition. In the first scenario, we have 7 iterations where we sample 200 subjects for each condition (*I* × *N_l_* = 7 × 200 = 1400). In the test condition, we then sub-sample these subjects into 10 studies (*K* = 10). In the second scenario, we double the amount of studies in the meta-analysis with *N_l_* = 400, *K* = 20 and *I* = 3. Finally, in the third scenario we include 35 studies in each meta-analysis which leads to *N_l_* = 700, *K* = 35 and *I* = 2. This is the maximum *K* as we then use all the subjects from the database while performing more than 1 iteration.

**Figure 2.**
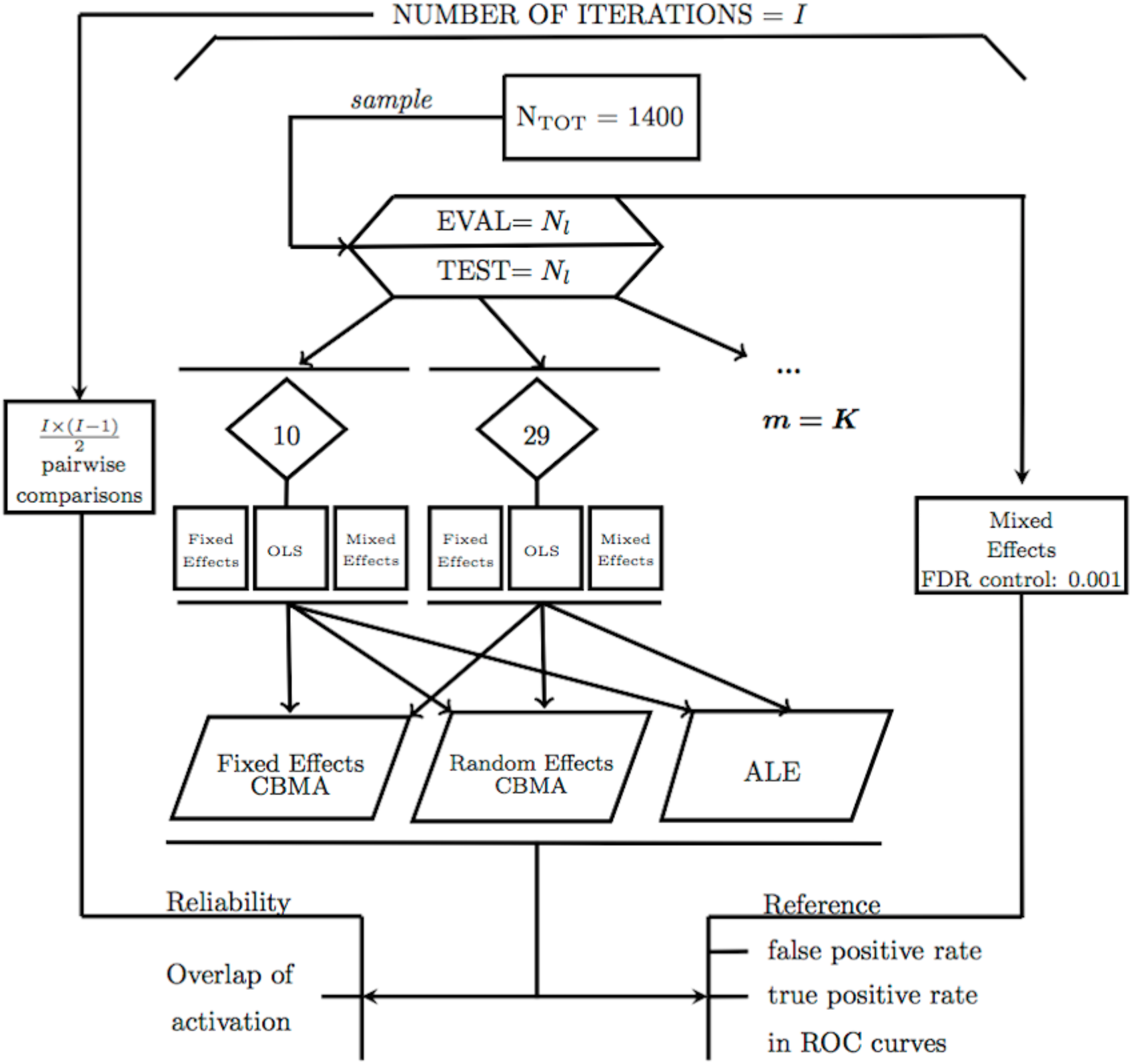
Design of the study illustrating the calculation of false positives and true positives and reliability using an evaluation condition (EVAL) and test condition (TEST).

#### 2.4.2 Test condition

The *K* studies in the test condition are all analysed using FSL, version 5.0.6. Every second level GLM model (FE, OLS and ME) is fitted to each of the *K* studies with the FLAME 1 + 2 option for the mixed effects models. We only test for average group activation.

To obtain local maxima, we search for clusters of significant activity in the *K* studies of the test condition. We choose this as clusters give an intuitive way of defining local maxima (i.e. the highest peak within each cluster). To control for multiple testing, we first determine a threshold such that the voxelwise false discovery rate (FDR) is controlled at level 0.05. Then, we determine clusters of significant voxels by using this FDR threshold as a cluster forming threshold in combination with a 26-point search algorithm. By doing so, we still obtain local maxima, but avoid clusterwise inference which is shown to be conservative (Eklund et al., 2016). The average observed cluster forming threshold in this study equals *Z* = 3.18. The resulting coordinates of the foci from each study with the number of subjects are then used as input for the ALE meta-analysis. The corresponding *t*-values (peak heights) are added for the fixed and random effects coordinate-based meta-analyses.

To identify significant voxels in the resulting meta-analyses, we apply the recommended procedures as described in section 2.3. For ALE, a voxelwise threshold uncorrected for multiple testing is used at level 0.001, as well as a cluster-level family-wise error (cFWE) correction for multiple testing at level 0.05. For the fixed and random effects CBMA we use a threshold at *Z* > 1 and at *P* = 0.005, uncorrected for multiple testing.

#### 2.4.3 Evaluation condition

Finally, the *N_l_* subjects in the evaluation condition are combined in one large, high powered study, using a mixed effects model. To control for multiple testing and balance sensitivity and specificity in this large sample, we apply a more conservative threshold such that the voxelwise FDR is controlled at level 0.001. The resulting map serves as a reference/benchmark image for the meta-analysis results obtained in the test condition. Note that a threshold for the sample in the evaluation condition could be chosen in different ways so deviations from the benchmark image should not be interpreted in an absolute manner but compared between methods in a relative manner. Furthermore, we do not model all available subjects into the evaluation condition, but a set of *N_l_* different subjects with respect to the test condition. This ensures that the evaluation condition is based on independent data. Next, by having an equal sample size in both conditions one can consider the evaluation condition as a perfect scenario in which all data is available for aggregation, while the test condition is the scenario in which we need to aggregate censored summary results in the form of peak coordinates.

### 2.5 Performance measures

To assess the performance of the different procedures for CBMA, we use two different measures: the balance between false positives and true positives in receiver operator characteristic (ROC) curves and activation reliability as a proxy for replicability.

#### 2.5.1 ROC curves

Statistical tests are often evaluated based on the extent to which they are able to minimize the number false positives (detecting signal where there is none) while maximizing the amount of true positive hits (detecting true signal). Receiver operator characteristic (ROC) curves plot the observed true positive rate (TPR) against the observed false positive rate (FPR) as the threshold for significance (*α*) is gradually incremented. To calculate true and false positives, we compare the results from the meta-analysis in the test condition with the reference image in the evaluation condition (EVAL on figure 2). The TPR or sensitivity is calculated as the number of voxels that are statistically significant in both the meta-analysis map and the reference map divided by the total number of voxels that is statistically significant in the reference map. The FPR or fall-out is calculated as the number of voxels that is statistically significant in the meta-analysis map but not in the reference map divided by the total number of voxels that is NOT statistically significant in the reference map.

Because the TPR and FPR are calculated voxelwise, we construct the ROC curves based on uncorrected *p* - values for the meta-analyses by incrementing the significance level, alpha, from 0 to 1. Finally, we average the *I* individual ROC curves and additionally use the area under the curve (AUC) as a summary measure. Higher AUC values indicate a better balance in discriminating between false positive and true positive voxels. We also plot the ROC calculate the AUC for that part of the curve for which *α* ∈ [0,0.1] by means of the standardized partial AUC (McClish, 1989).

Since the ALE algorithm uses an MNI brain template with a higher resolution (2 mm voxels, dimensions 91 × 109 × 91) than the (pre-processed) IMAGEN data (3 mm voxels, dimensions 53 × 63 × 46), the reference image is also resampled to a higher resolution so that it matches the resolution of the ALE images. We apply a linear affine transformation with 12 degrees of freedom from the EPI template of the IMAGEN dataset to the MNI brain template, using a correlation ratio cost function (Jenkinson et al., 2002) and trilinear interpolation in FSL. As the fixed and random effects meta-analyses model the local maxima using the same brain template as the IMAGEN data, no such transformation is needed here to calculate the ROC curves.

#### 2.5.2 Reliability

We consider activation reliability as an indicator for the success of replicating results. We define replicability as the ability to repeat the results of an experiment using the exact same materials, procedures and methods, but with a different set of subjects. There is no consensus in the literature on this definition as other authors use terms such as strong replicable results or direct reproduction to indicate the same concept (Patil et al., 2016; Pernet and Poline, 2015). We quantify reliability in two ways.

First, we measure the overlap of results between iterations of the same analysis pipeline. We calculate the percent overlap of activation (Maitra, 2010) between all 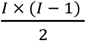 pairwise combinations of the *I* unique iterations of the design (figure 2). Let *V_a,b_* represent the intersection of statistically significant voxels in image *a* and *b*, *V_a_* the amount of statistically significant voxels in image *a* and *V_b_* the amount of statistically significant voxels in image *b*. The overlap *ω_a,b_* is then defined as:

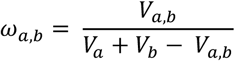

This measure ranges from 0 (no overlap) to 1 (perfect overlap). Note that this is an adaptation of the Dice (1945) or the Sørensen (1948) similarity coefficient.

As a second method to quantify reliability, we describe the amount of unique information captured in each iteration. We first quantify the number of times out of the *I* iterations a voxel is declared significant and visualize this on a heatmap. We do the same for the *I* reference images from the evaluation condition. As a comparison, we include the average effect size map obtained using again the reference images.

Next, we run a 26 point search clustering algorithm on each thresholded meta-analysis to calculate the frequency of clusters of at least one statistically significant voxel. We record the average cluster size expressed in number of voxels. We then assess the number of unique clusters across the pairwise combinations. A cluster of statistically significant voxels in image *a* is unique if no single voxel from this cluster overlaps with a cluster of statistically significant voxels in the paired image *b*. We finally determine the amount of these unique clusters that are large (we have set the threshold for large at 50 voxels) and divide this by the total amount of statistically significant clusters to obtain the proportion of large unique clusters. Additionally, we study the number of clusters and cluster sizes for both unique and overlapping clusters to get an overview, independent of the chosen threshold on the cluster size. Given a sample size, smaller amounts of (large) unique clusters imply a higher pairwise reliability.

## 3 Results

### 3.1 ROC curves

In figure 3, 5 and 7 we present the average ROC curves (over iterations) that show the observed true positive rate against the observed false positive rate for *K* = 10, 20 and 35 over the entire range of *α*. In figure 4, 6 and 8 we present the average ROC curves for K = 10, 20 and 35 when *α* ∈ [0,0.1]. To condense this section, we only discuss results based on the entire range of *α*. We observe the same patterns emerging when *α* ∈ [0,0.1]. The overall AUC is high, but recall that given that comparisons are made with the reference image, all values should be used for relative comparisons as the absolute AUC will depend on how the reference image is determined.

**Figure 3:**
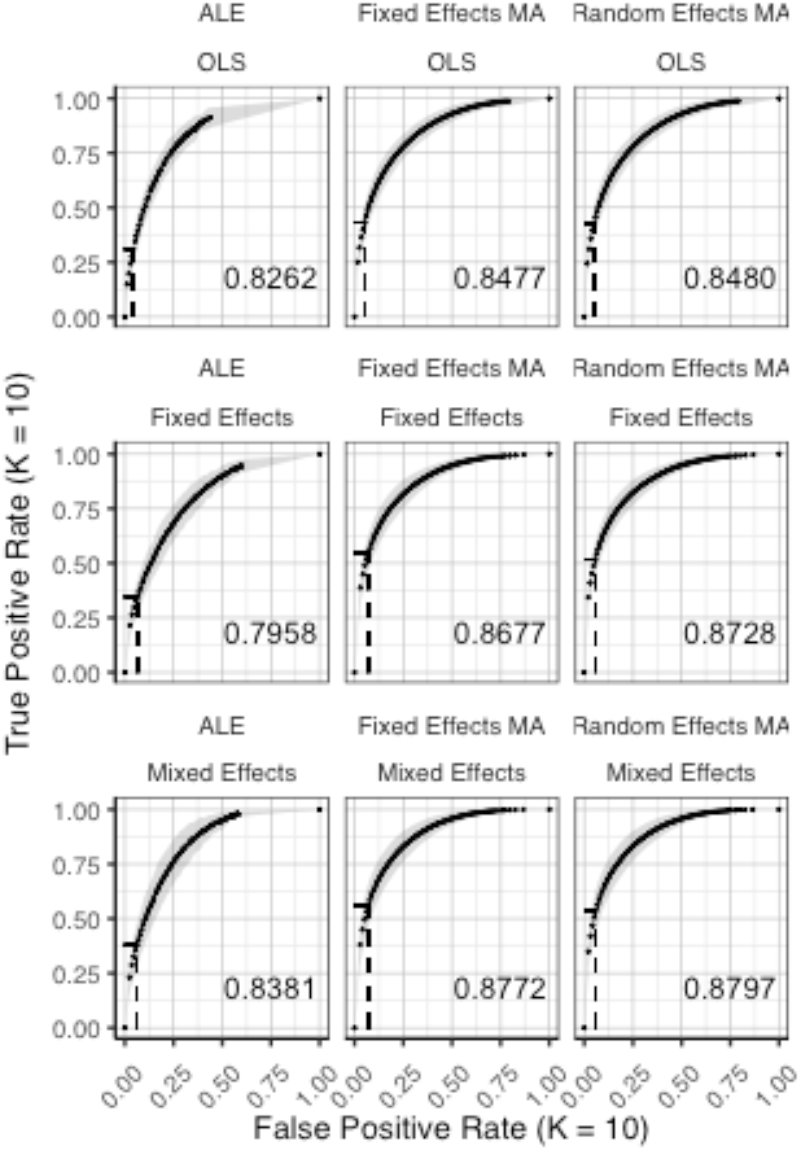
complete ROC curves (± 1 standard deviation), averaged over *I* = 7 iterations plotting the observed true positive rate against the observed false positive rate for *K* = 10. The columns correspond to the coordinate-based meta-analyses (left: ALE uncorrected procedure, middle: fixed effects meta-analysis, right: random effects meta-analysis). The rows correspond to the second level GLM pooling models (top: OLS, middle: fixed effects, bottom: mixed effects). For each of those, the area under the curve (AUC) is calculated and shown within the plot. The drop-down lines correspond to the point at which the pre-specified nominal level is set at an uncorrected *α* level of 0.05.

**Figure 4:**
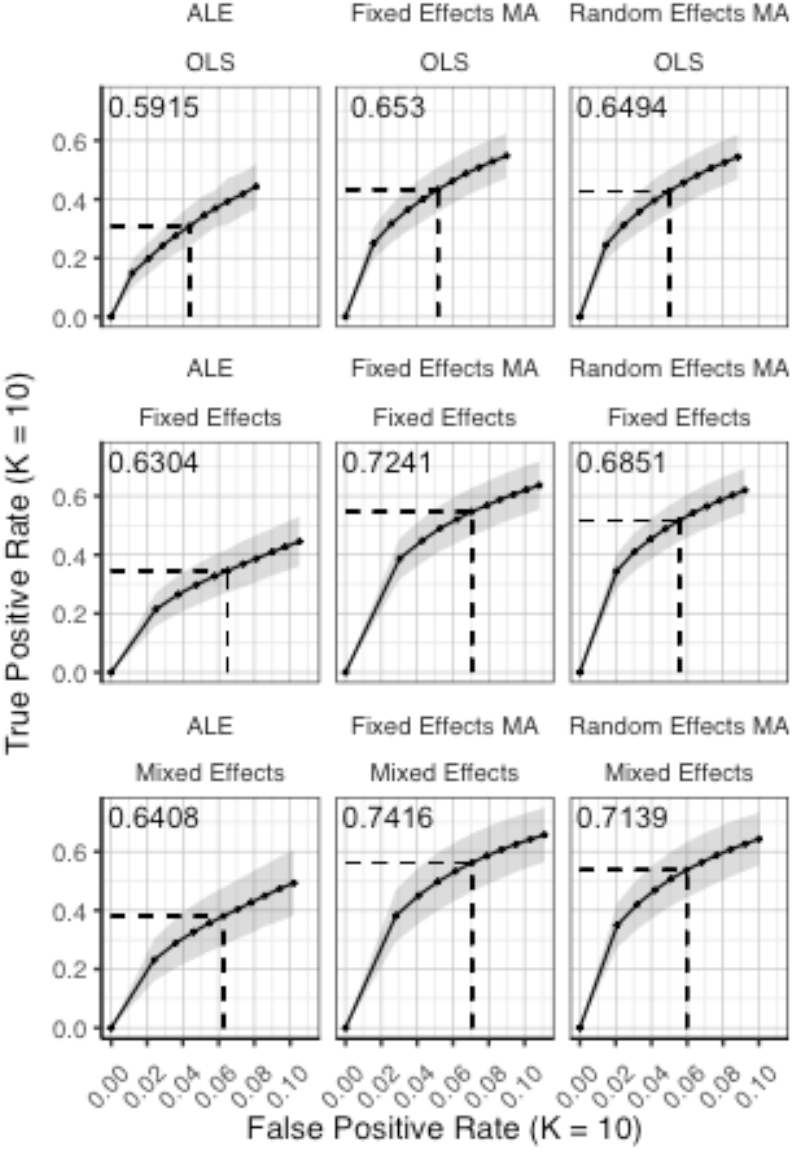
identical ROC curves as in figure 3 (*K* = 10), but only for *α* ∈ [0,0.1]. The area under the curve is calculated through a standardized partial AUC. The columns correspond to the coordinate-based meta-analyses (left: ALE uncorrected procedure, middle: fixed effects meta-analysis, right: random effects meta-analysis). The rows correspond to the second level GLM pooling models (top: OLS, middle: fixed effects, bottom: mixed effects). The drop-down lines correspond to the point at which the pre-specified nominal level is set at an uncorrected *α* level of 0.05.

**Figure 5.**
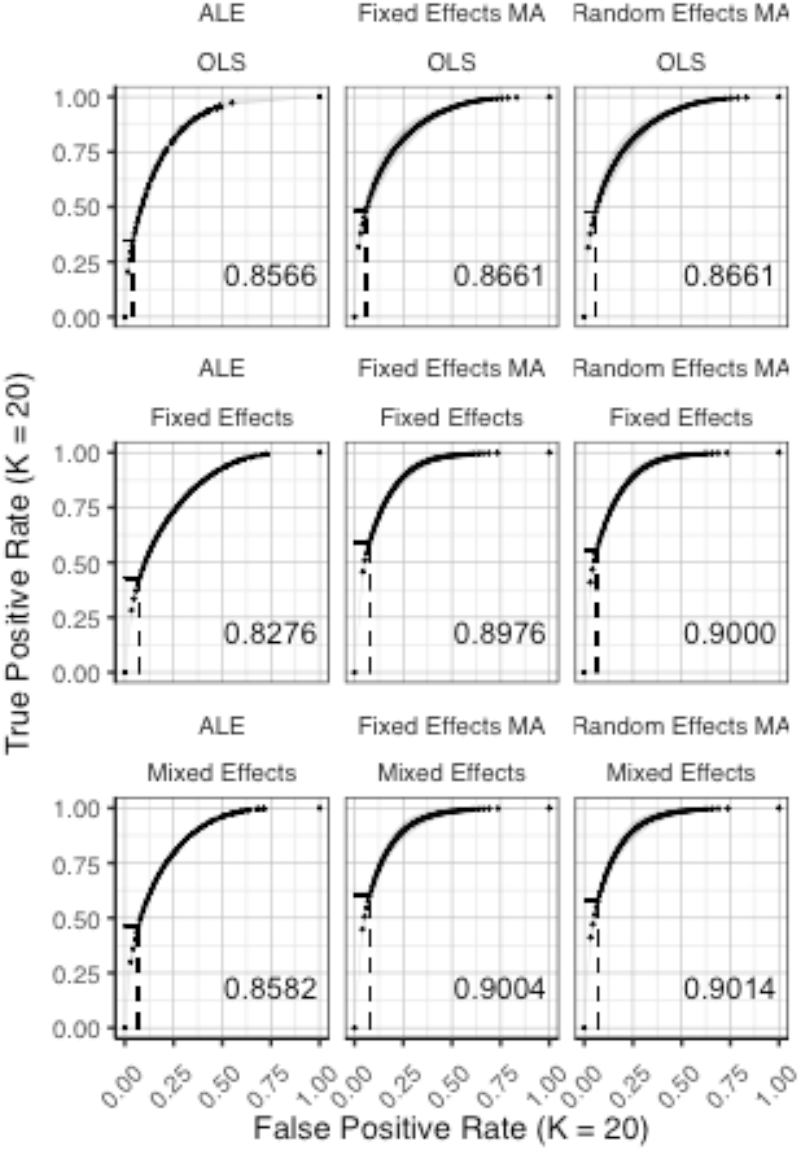
ROC curves (± 1 standard deviation), averaged over *I* = 3 iterations plotting the observed true positive rate against the observed false positive rate for *K*=20. The columns correspond to the coordinate-based meta-analyses (left: ALE uncorrected procedure, middle: fixed effects meta-analysis, right: random effects metaanalysis). The rows correspond to the second level GLM pooling models (top: OLS, middle: fixed effects, bottom: mixed effects). For each of those, the area under the curve (AUC) is calculated and shown within the plot. The drop-down lines correspond to the point at which the pre-specified nominal level is set at an uncorrected *α* level of 0.05.

**Figure 6.**
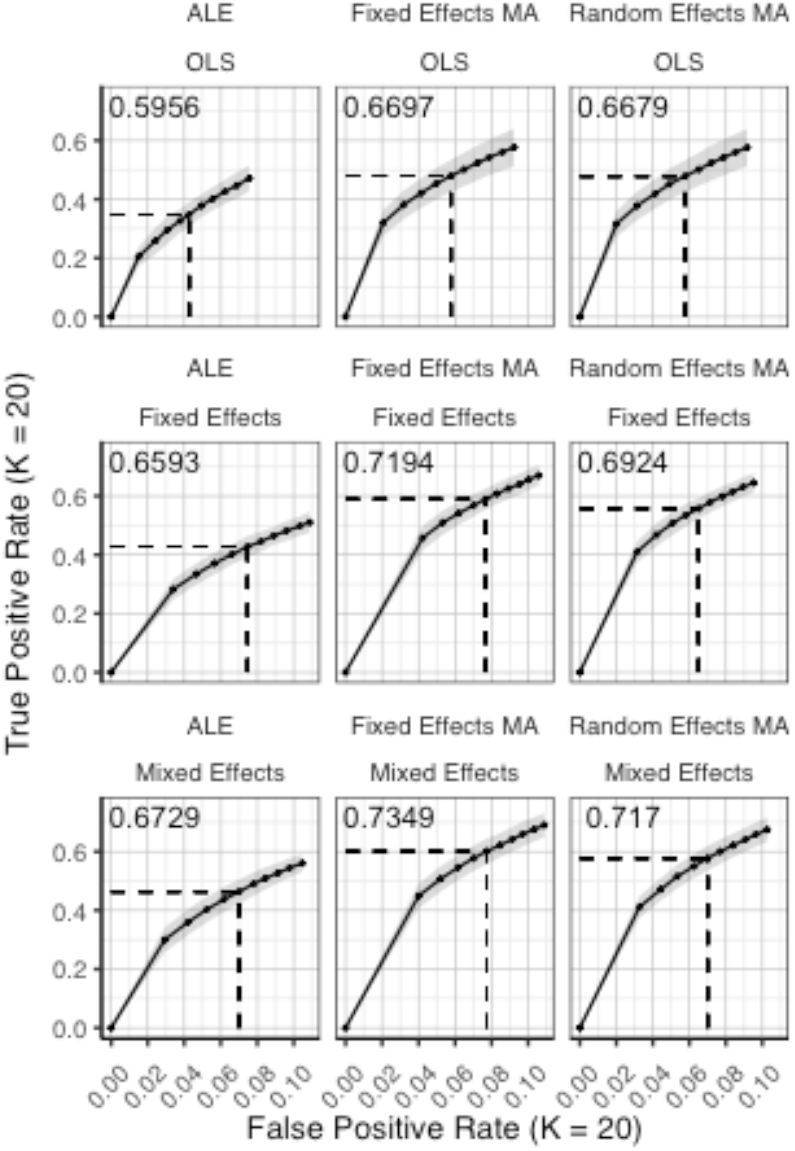
identical ROC curves as in figure 5 (*K* = 20), but only for *α* ∈ [0,0.1]. The area under the curve is calculated through a standardized partial AUC. The columns correspond to the coordinate-based meta-analyses (left: ALE uncorrected procedure, middle: fixed effects meta-analysis, right: random effects meta-analysis). The rows correspond to the second level GLM pooling models (top: OLS, middle: fixed effects, bottom: mixed effects). The drop-down lines correspond to the point at which the pre-specified nominal level is set at an uncorrected *α* level of 0.05.

**Figure 7.**
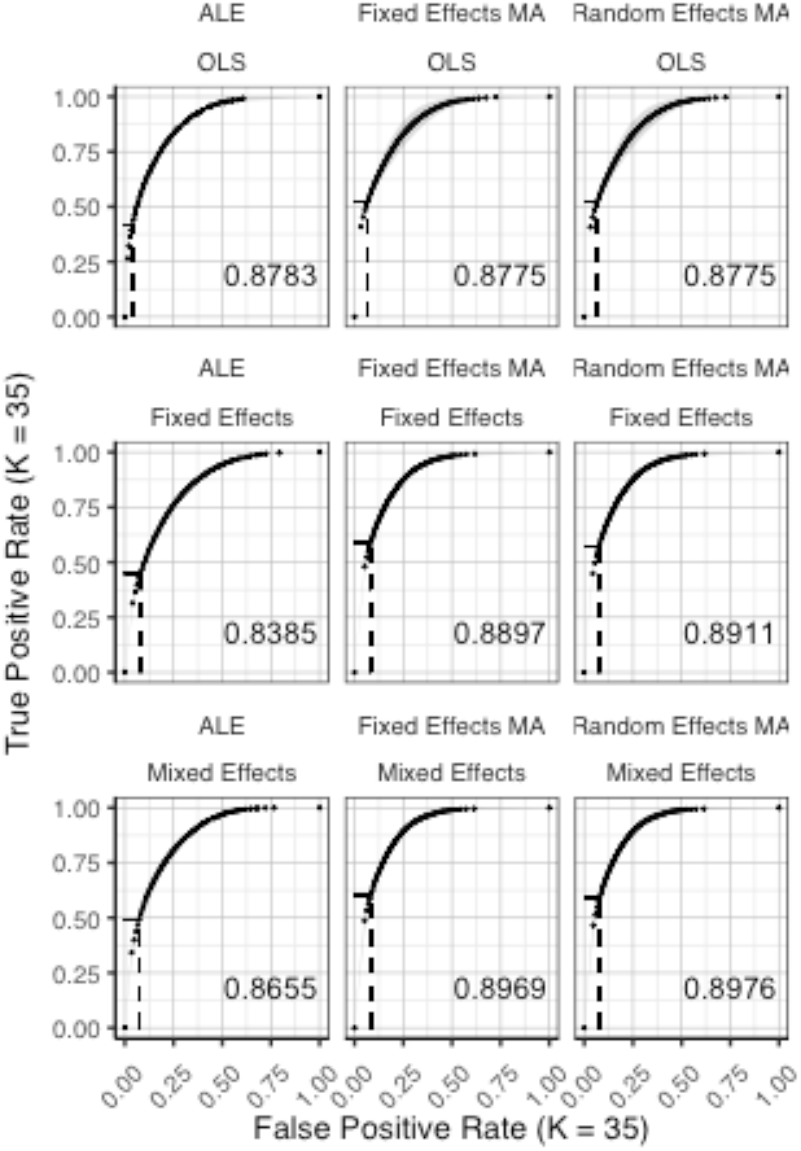
ROC curves (*±* 1 standard deviation), averaged over *I* = 2 iterations plotting the observed true positive rate against the observed false positive rate for *K*=35. The columns correspond to the coordinate-based meta-analyses (left: ALE uncorrected procedure, middle: fixed effects meta-analysis, right: random effects metaanalysis). The rows correspond to the second level GLM pooling models (top: OLS, middle: fixed effects, bottom: mixed effects). For each of those, the area under the curve (AUC) is calculated and shown within the plot. For each of those, the area under the curve (AUC) is calculated and shown within the plot. The drop-down lines correspond to the point at which the pre-specified nominal level is set at an uncorrected *α* level of 0.05.

**Figure 8.**
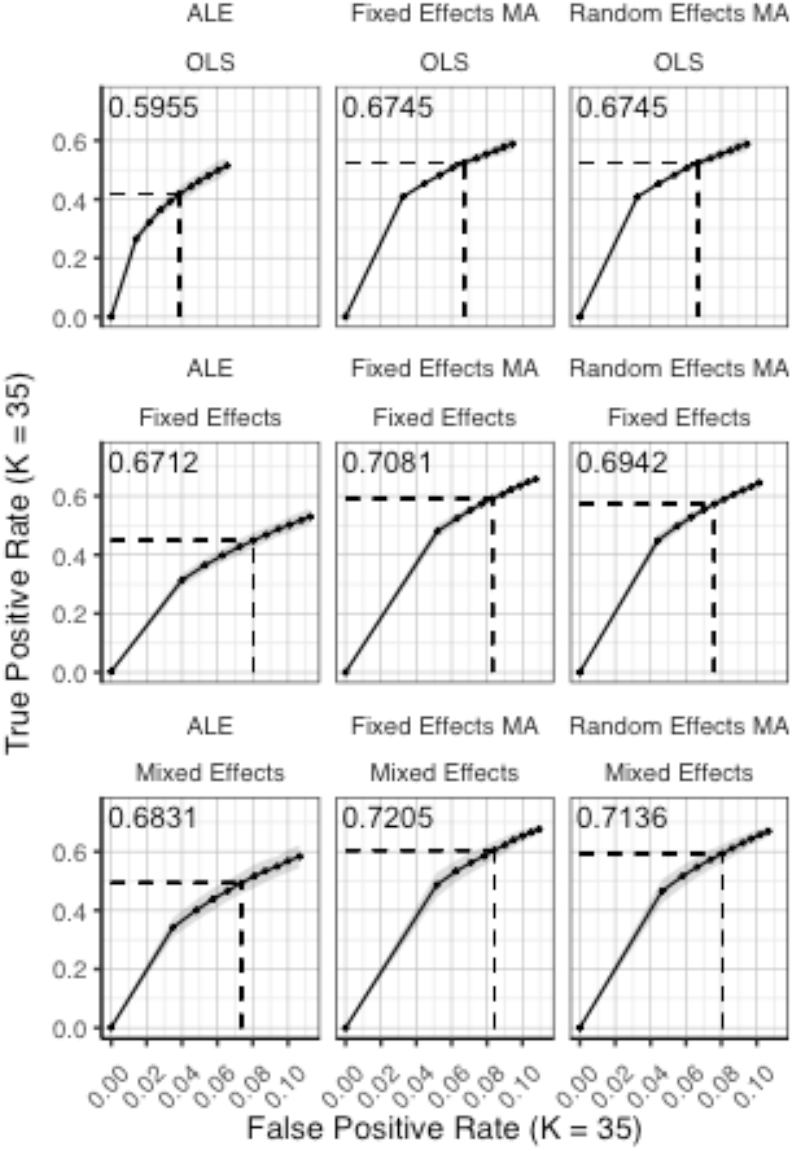
identical ROC curves as in figure 7 (*K* = 35), but only for *α* ∈ [0,0.1]. The area under the curve is calculated through a standardized partial AUC. The columns correspond to the coordinate-based meta-analyses (left: ALE uncorrected procedure, middle: fixed effects meta-analysis, right: random effects meta-analysis). The rows correspond to the second level GLM pooling models (top: OLS, middle: fixed effects, bottom: mixed effects). The drop-down lines correspond to the point at which the pre-specified nominal level is set at an uncorrected *α* level of 0.05.

We observe higher AUC values using fixed and random effects models compared to ALE. The only exception is observed for the combination of OLS and ALE for *K* = 35. Small differences are observed between the fixed and random effects meta-analysis with generally higher AUC values for random effects meta-analyses. The observed TPR at an uncorrected threshold of 0.05 never exceeds 0.5 for ALE in any of the scenarios, while the TPR of the fixed and random effects CBMA methods approaches 0.6 when combining mixed or fixed group level models with a higher amount of studies in the meta-analysis.

The AUC reveals an interaction between group level models and CBMA methods. In the fixed and random effects meta-analysis an OLS model is associated with lower values of the AUC compared to fixed and mixed effects models, regardless of the amount of studies in the meta-analysis. While the difference between fixed and mixed effects group models is minimal, the mixed effects model consistently outperforms the fixed effects model.

When using the ALE algorithm however, the lowest AUC is consistently associated with a fixed effects group model for all study set sizes K. Only with the highest amount of studies in the meta-analysis (*K* = 35), does an OLS group model outperform the mixed effects model. The combination of an OLS model with the ALE algorithm not only leads to a lower observed TPR at an uncorrected threshold of 0.05, but also a lower observed FPR.

Finally, for all CBMA methods, increasing the number of studies in the meta-analysis from 10 to 20 results in a higher AUC. The average AUC of the meta-analyses, regardless of the group level models, increases for *K* = 10 from 0.82 (ALE), 0.86 (fixed effects MA) and 0.87 (random effects MA) to respectively 0.85, 0.89 and 0.89 in *K* = 20. Adding even more studies (*K* = 35) is associated with a further increase to 0.86 of the average AUC for ALE, but not for the fixed (0.89) and random effects (0.89) meta-analyses.

Overall, the best balance between TPR and FPR detection is observed when using mixed effects group level models together with random effects meta-analyses.

### 3.2 Reliability

Figures 9 and 10 display the percent overlap of activation for *K* = 10, 20 and 35. Noticeably, the overlap values have a wide range from 0.07 (OLS, ALE cFWE, *K* = 10) to a moderate 0.69 (fixed effects group level model, random effects MA, *K* = 35). Average overlap values over *I* iterations and the group level models/CBMA methods can be found in table 1. Again, as the overlap between thresholded maps depends on the chosen threshold, it is better to focus on the relative performances of the group level models and methods for CBMA.

**Figure 9.**
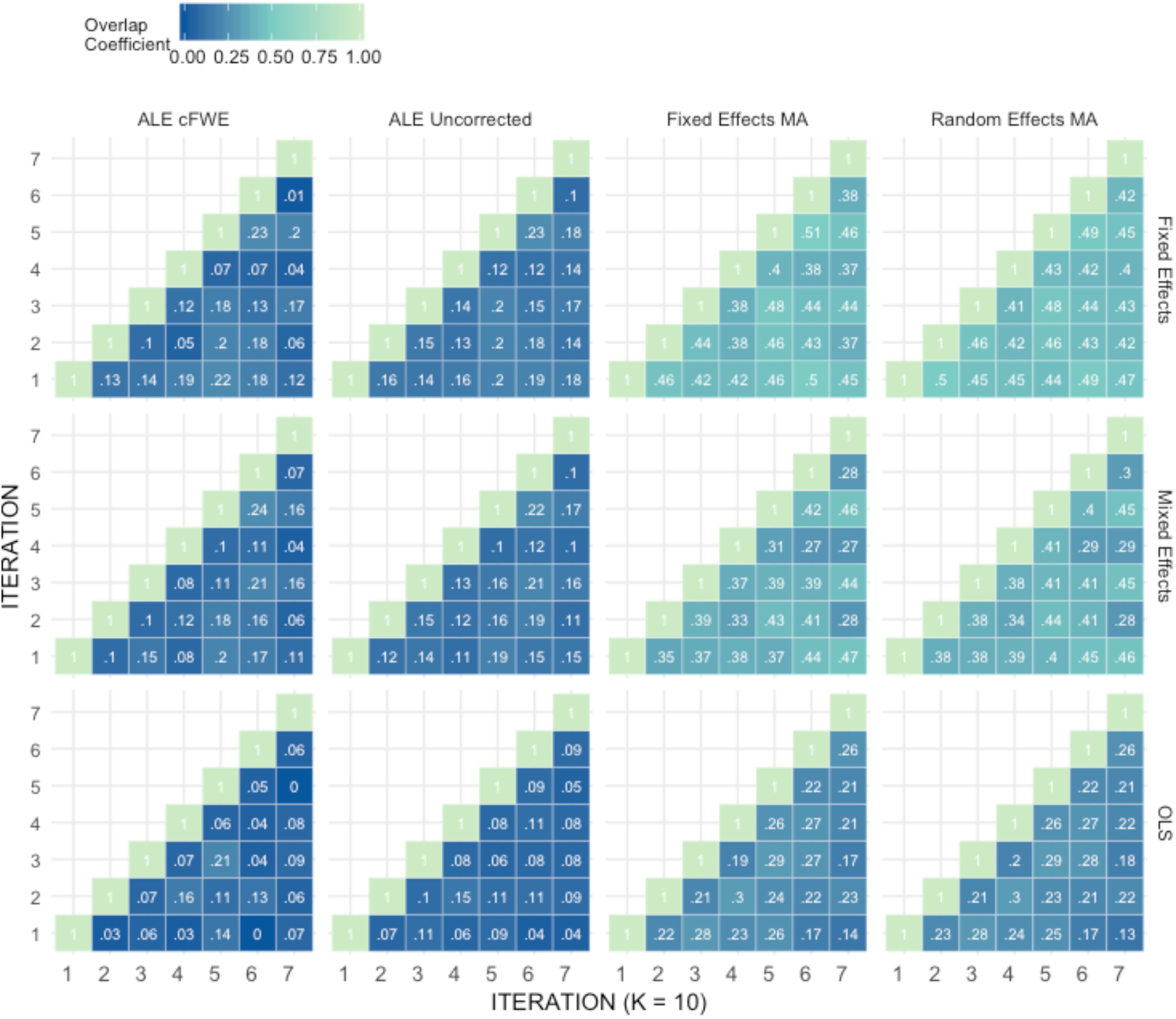
Percent overlap of activation (*ω_a,b_*) from all pairwise comparisons for *K* = 10. The rows represent the group level models (top to bottom: fixed effects, mixed effects and OLS). The columns represent the thresholded meta-analyses. From left to right: ALE cFWE at 0.05, ALE uncorrected at 0.001 and fixed and random effects CBMA at 0.005 with *Z* > 1.

**Figure 10.**
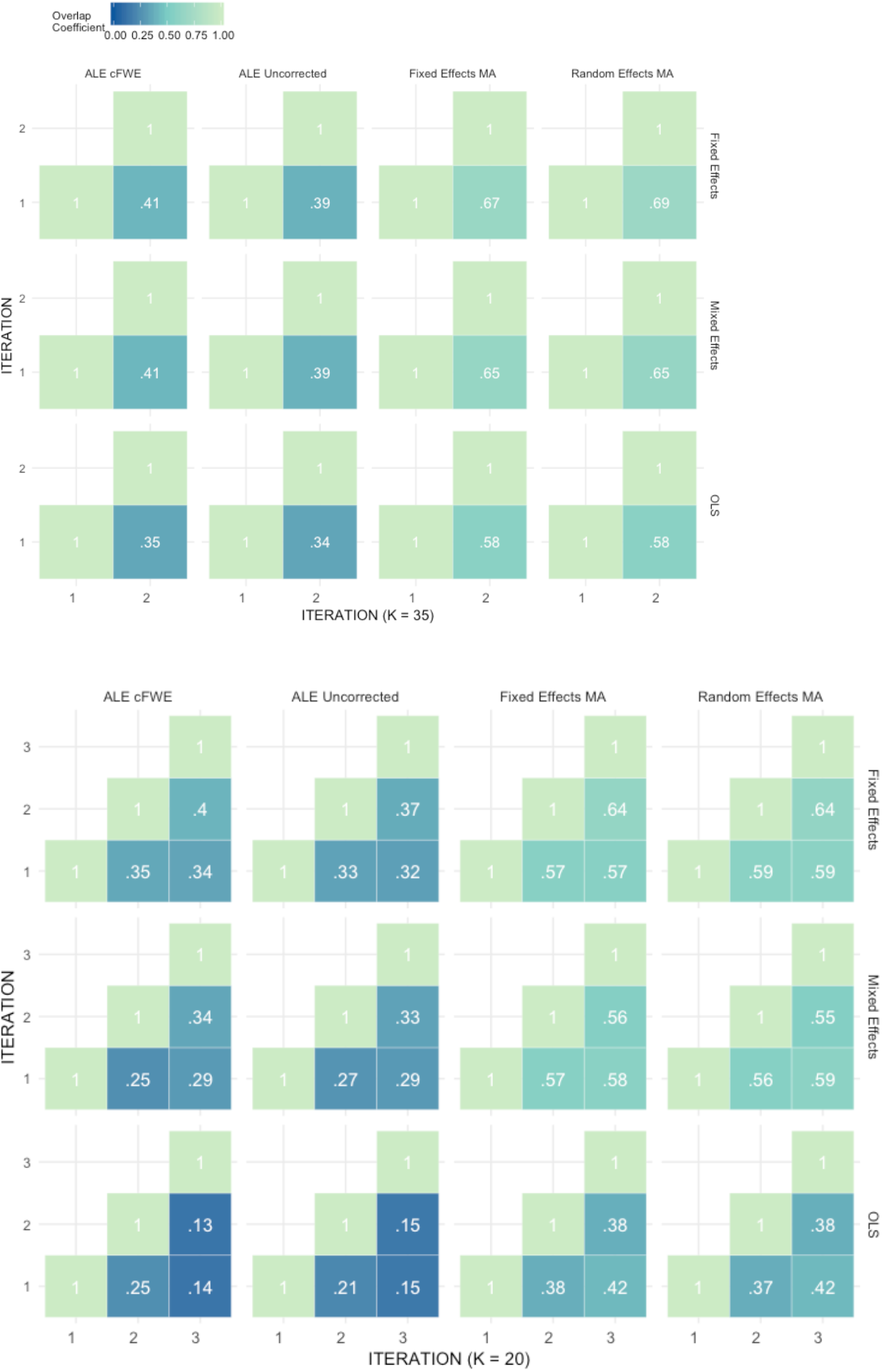
Percent overlap of activation (*ω_a,b_*) from all pairwise comparisons for K = 20 (bottom) and 35 (top). The rows represent the group level models (top to bottom: fixed effects, mixed effects and OLS). The columns represent the thresholded meta-analyses. From left to right: ALE cFWE at 0.05, ALE uncorrected at 0.001 and fixed and random effects CBMA at 0.005 with *Z* > 1.

**Table 1.** Averaged overlap values over the *I* iterations and the CBMA methods (top) and over the *I* iterations and the group level models (bottom) for each *K*.

| Average overlap over $I$ and CBMA methods | | | |
| --- | --- | --- | --- |
| K | Fixed effects | Mixed effects | OLS |
| 10 | 0.29 | 0.26 | 0.15 |
| 20 | 0.48 | 0.43 | 0.28 |
| 35 | 0.54 | 0.52 | 0.46 |

| Average overlap over $I$ and group level models | | | | |
| --- | --- | --- | --- | --- |
| K | Fixed Effects MA | Random Effects MA | ALE Uncorrected | ALE cFWE |
| 10 | 0.34 | 0.35 | 0.13 | 0.11 |
| 20 | 0.52 | 0.52 | 0.27 | 0.28 |
| 35 | 0.64 | 0.64 | 0.38 | 0.39 |

Similar to the ROC curves, we observe higher overlap when more studies are added to the meta-analysis. Furthermore, both ALE thresholding methods are associated with lower values of overlap compared to the fixed and random effects meta-analysis. In contrast to the ROC curves, the maximum overlap value observed in ALE is low and does not approach the performance of the fixed and random effects meta-analysis. We only observe small differences between the fixed and random effects meta-analysis. For *K* = 10, we observe mostly higher values using a random effects meta-analysis.

Regarding the group level models, OLS models are associated with lower coefficients of overlap than fixed and mixed effects models. In general, we observe higher values using fixed effects models compared to mixed effects models, though these differences are much smaller. These patterns are similar regardless of the CBMA method and study set size *K*.

Given the results on the overlap values, we look for similar patterns using the heatmaps at MNI z-coordinate 50 for *K* = 10 (left part of figure 11.A), *K* = 20 (right part of figure 11.A) and *K* = 35 (figure 12.A) and in the results detailing the amount of unique information in each iteration (table 2).

**Figure 11.**
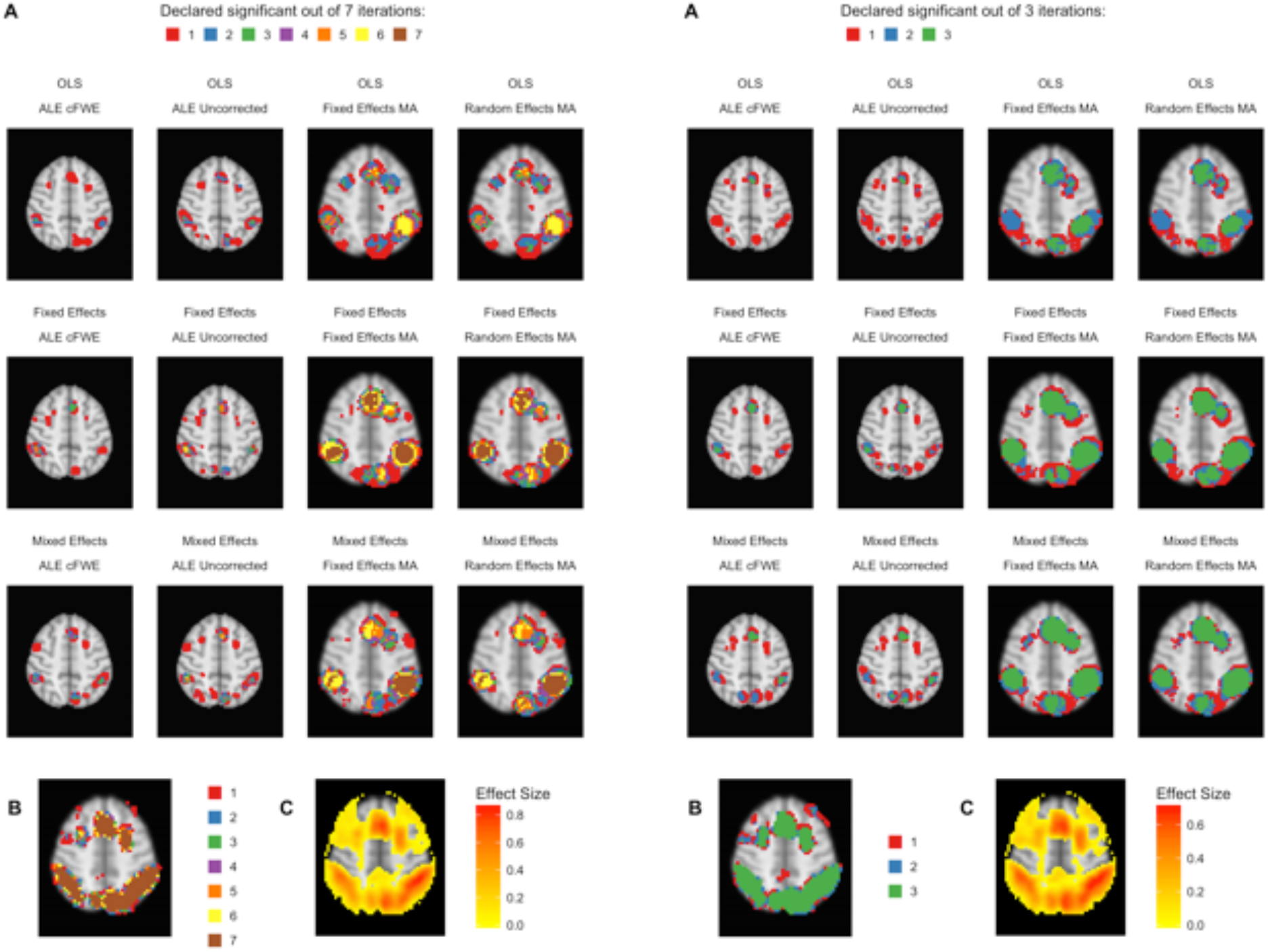
Heatmaps of MNI *z*-coordinate 50 for *K* = 10 (left) and *K* = 20 (right). A: the number of iterations in which each voxel has been declared statistically significant for each combination of a group level model (row-wise) and thresholded meta-analysis (column-wise). B: the number of iterations in which each voxel of the reference images has been declared statistically significant. Areas of interest involve the supramarginal gyrus (posterior division), superior parietal lobule and angular gyrus. C: average effect size of the reference images over the iterations.

**Figure 12.**
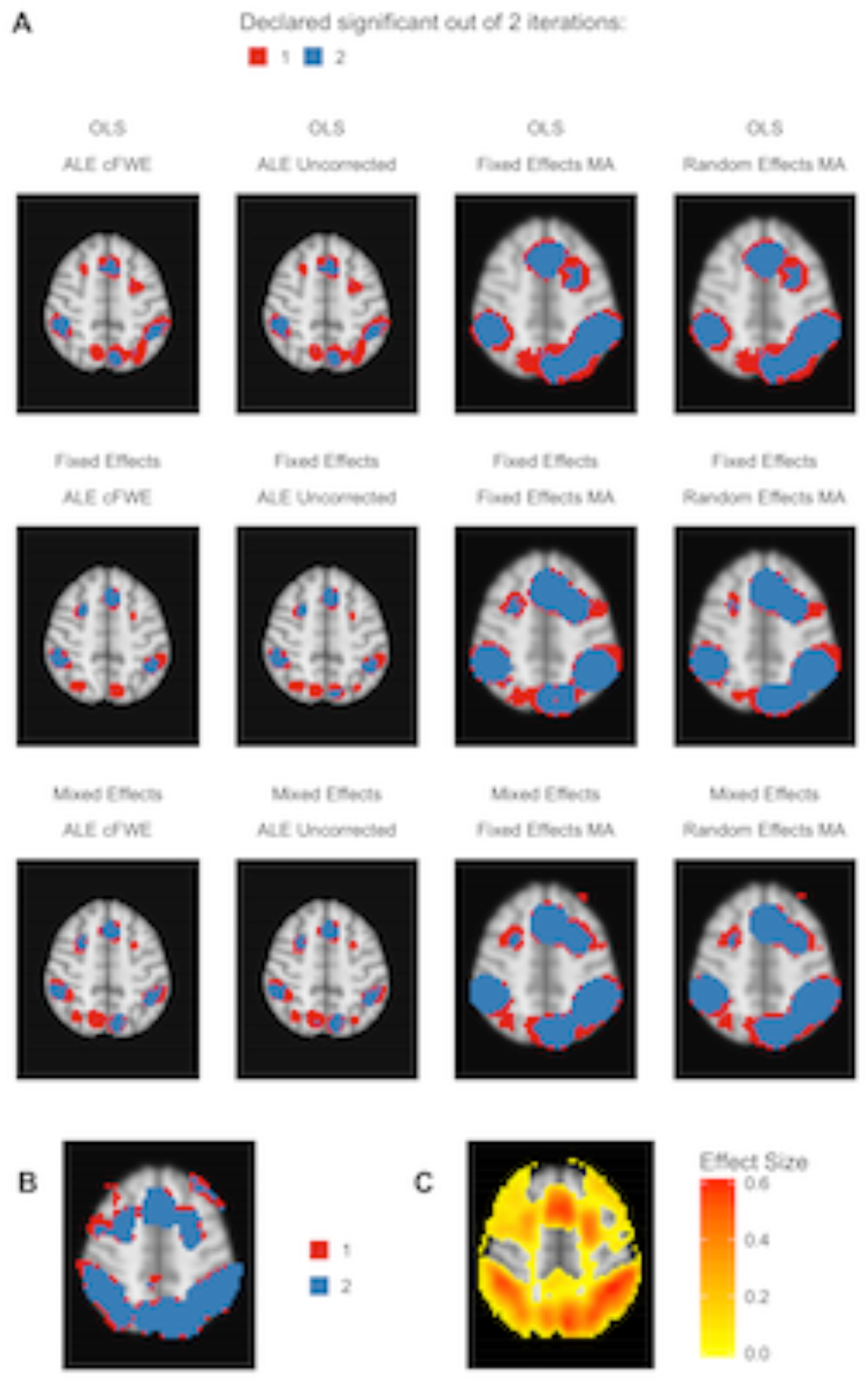
Heatmap of MNIz-coordinate 50 for *K* = 35. A: the number of iterations in which each voxel has been declared statistically significant for each combination of a group level model (row-wise) and thresholded meta-analysis (column-wise). B: the number of iterations in which each voxel of the reference images has been declared statistically significant. Areas of interest involve the supramarginal gyrus (posterior division), superior parietal lobule and angular gyrus. C: average effect size of the reference images over the iterations.

**Table 2.** Descriptive results of the thresholded meta-analyses in a replication setting. For each study set size (*K*), *I* replicated images are compared pairwise. Shown in the table are the averages (over *I*) of the amount of clusters and the size of these clusters. Next to it are the averages (over 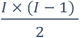 pairwise comparisons) of the amount of clusters that are unique to one of the paired comparisons, the amount of large (i.e. more than 50 voxels) unique clusters and the percentage of the total amount of clusters that are large unique clusters.

| K | Group Model | Meta-analysis | I | Amount of clusters |  | Voxels in clusters |  | Unique clusters |  | Large uni. clust. |  | Percentage |
| --- | --- | --- | --- | --- | --- | --- | --- | --- | --- | --- | --- | --- |
|  |  |  |  | mean | sd | mean | sd | mean | sd | mean | sd | large clusters |
| 10 | Fixed Effects | Fixed Effects MA | 7 | 23.1 | 6.1 | 206.7 | 71.4 | 11.67 | 5.83 | 2.00 | 1.48 | 0.09 |
|  |  | Random Effects MA | 7 | 19.6 | 3.1 | 186.4 | 43.9 | 9.33 | 2.88 | 1.45 | 1.52 | 0.08 |
|  |  | ALE Uncorrected | 7 | 50.3 | 7.2 | 53.1 | 11.1 | 31.10 | 6.66 | 4.14 | 2.72 | 0.08 |
|  |  | ALE cFWE | 7 | 11.0 | 1.3 | 155.0 | 31.8 | 5.71 | 1.94 | 5.71 | 1.94 | 0.52 |
|  | OLS | Fixed Effects MA | 7 | 19.9 | 6.1 | 132.1 | 59.6 | 11.86 | 4.95 | 2.69 | 2.41 | 0.14 |
|  |  | Random Effects MA | 7 | 20.6 | 7.4 | 126.6 | 68.6 | 12.43 | 5.92 | 2.62 | 2.23 | 0.13 |
|  |  | ALE Uncorrected | 7 | 31.9 | 6.8 | 41.7 | 15.2 | 22.95 | 5.65 | 3.17 | 2.04 | 0.10 |
|  |  | ALE cFWE | 7 | 4.9 | 2.9 | 136.5 | 49.0 | 3.10 | 2.35 | 3.10 | 2.35 | 0.63 |
|  | Mixed Effects | Fixed Effects MA | 7 | 22.9 | 4.7 | 189.0 | 50.8 | 12.19 | 3.98 | 2.36 | 1.41 | 0.10 |
|  |  | Random Effects MA | 7 | 21.4 | 3.4 | 169.2 | 44.0 | 10.38 | 3.22 | 1.40 | 0.94 | 0.07 |
|  |  | ALE Uncorrected | 7 | 49.1 | 8.1 | 54.0 | 14.1 | 30.57 | 7.03 | 4.26 | 2.31 | 0.09 |
|  |  | ALE cFWE | 7 | 11.6 | 1.5 | 147.3 | 26.1 | 5.95 | 1.77 | 5.95 | 1.77 | 0.52 |
| 20 | Fixed Effects | Fixed Effects MA | 3 | 19.3 | 5.1 | 438.1 | 145.5 | 8.00 | 4.29 | 2.17 | 1.83 | 0.12 |
|  |  | Random Effects MA | 3 | 17.0 | 1.7 | 394.8 | 71.7 | 4.67 | 1.86 | 0.50 | 0.55 | 0.03 |
|  |  | ALE Uncorrected | 3 | 52.3 | 8.4 | 128.1 | 42.0 | 22.67 | 7.37 | 4.67 | 2.16 | 0.09 |
|  |  | ALE cFWE | 3 | 21.0 | 2.6 | 264.8 | 53.4 | 3.33 | 2.07 | 3.33 | 2.07 | 0.16 |
|  | OLS | Fixed Effects MA | 3 | 21.3 | 8.7 | 248.9 | 105.2 | 12.33 | 7.55 | 3.17 | 0.75 | 0.15 |
|  |  | Random Effects MA | 3 | 20.3 | 8.4 | 251.0 | 97.0 | 11.00 | 7.40 | 3.00 | 0.63 | 0.15 |
|  |  | ALE Uncorrected | 3 | 47.0 | 11.1 | 64.9 | 17.3 | 29.00 | 9.72 | 5.17 | 2.14 | 0.11 |
|  |  | ALE cFWE | 3 | 12.3 | 1.5 | 181.6 | 36.9 | 5.33 | 1.97 | 5.33 | 1.97 | 0.44 |
|  | Mixed Effects | Fixed Effects MA | 3 | 20.7 | 4.5 | 389.8 | 122.1 | 8.67 | 4.27 | 1.00 | 1.10 | 0.05 |
|  |  | Random Effects MA | 3 | 21.0 | 1.0 | 318.6 | 44.0 | 9.00 | 0.89 | 1.00 | 0.89 | 0.05 |
|  |  | ALE Uncorrected | 3 | 50.7 | 6.1 | 123.9 | 36.5 | 26.67 | 5.16 | 5.67 | 2.80 | 0.11 |
|  |  | ALE cFWE | 3 | 18.3 | 2.5 | 279.4 | 54.4 | 5.33 | 2.07 | 5.33 | 2.07 | 0.29 |
| 35 | Fixed Effects | Fixed Effects MA | 2 | 14.50 | 2.12 | 735.33 | 193.06 | 9.50 | 2.12 | 3.50 | 2.12 | 0.25 |
|  |  | Random Effects MA | 2 | 12.50 | 3.54 | 793.10 | 308.16 | 4.50 | 3.54 | 2.00 | 1.41 | 0.17 |
|  |  | ALE Uncorrected | 2 | 54.50 | 0.71 | 182.37 | 5.91 | 21.50 | 0.71 | 6.50 | 4.95 | 0.12 |
|  |  | ALE cFWE | 2 | 25.50 | 0.71 | 347.01 | 37.16 | 4.50 | 0.71 | 4.50 | 0.71 | 0.17 |
|  | OLS | Fixed Effects MA | 2 | 14.00 | 2.83 | 587.50 | 167.23 | 7.00 | 2.83 | 1.50 | 0.71 | 0.11 |
|  |  | Random Effects MA | 2 | 13.50 | 2.12 | 600.95 | 144.08 | 6.50 | 2.12 | 1.50 | 0.71 | 0.11 |
|  |  | ALE Uncorrected | 2 | 41.00 | 8.49 | 148.97 | 31.85 | 20.00 | 8.49 | 4.00 | 0.00 | 0.10 |
|  |  | ALE cFWE | 2 | 15.50 | 0.71 | 350.41 | 9.69 | 3.50 | 0.71 | 3.50 | 0.71 | 0.22 |
|  | Mixed Effects | Fixed Effects MA | 2 | 19.50 | 4.95 | 566.47 | 229.15 | 12.50 | 4.95 | 3.50 | 3.54 | 0.18 |
|  |  | Random Effects MA | 2 | 17.50 | 3.54 | 578.73 | 203.84 | 9.50 | 3.54 | 3.00 | 2.83 | 0.17 |
|  |  | ALE Uncorrected | 2 | 56.00 | 7.07 | 182.71 | 45.69 | 22.00 | 7.07 | 5.50 | 0.71 | 0.10 |
|  |  | ALE cFWE | 2 | 22.00 | 2.83 | 402.95 | 27.23 | 5.00 | 2.83 | 5.00 | 2.83 | 0.23 |

Regarding ALE, we clearly observe smaller regions of activation with a higher percentage of large unique clusters compared to the fixed and random effects meta-analysis, especially in small *K*. However, we do observe convergence in the ALE results to the brain regions characterized by (1) consistent statistically significant declared voxels (panel B in figure 11 and 12) and (2) high effect sizes in the reference images (panel C in figure 11 and 12). The fixed and random effects meta-analyses do detect larger regions, but are not necessarily constrained to the exact spatial shape of activated regions observed in the reference images.

The difference in the degree of unique information between uncorrected ALE and ALE cFWE is more detailed than the observed overlap values. Uncorrected ALE is associated with the highest (out of any meta-analysis) detection rate of small clusters. This in turn leads to an inflated number of (small and large) unique clusters. However, we observe the highest percentages of large unique clusters using ALE cFWE. Only small differences between the fixed and random effects meta-analyses are observed.

Regarding the group level models, we see on average less and smaller clusters of statistically significant voxels associated with the OLS group level models compared to the fixed and mixed effects models. This is true for every study set size *K*. However, for small study set sizes such as *K* = 10 and 20, the OLS model is associated with a higher percentage of large unique clusters. For *K*= 35, this is the opposite as the OLS model has on average the lowest percentage of large unique clusters. The fixed and mixed effects group level models show in most cases similar values. We include the distributions of the number of overlapping and unique detected clusters as well as the cluster sizes in the appendix. These distributions show the same patterns as depicted in table 2.

To conclude, models such as the OLS group level model (for *K* = 10 and 20) and the ALE meta-analyses that are characterized with low overlap values are either associated with smaller clusters of statistically significant voxels or higher percentages of large unique clusters.

### 3.3 Between study variability

We observe no substantial differences between the fixed and random effects meta-analysis in most results. Since we are working with one large database of a homogenous sample executing the same paradigm, between study variability is limited.

To investigate this further, we look at the between study variability, estimated by *τ*^2^ in the weights (U_*νm*_ equation (11)) of the random-effects meta-analysis for *K* = 10. In figure 13, we display the average *t*-map (over 7 iterations) of the reference images over 4 slices along the z-axis. We then plot the estimated *τ*^2^ from the random effects meta-analyses combined with the statistically significant voxels depicting the weighted averages of the random effects meta-analysis.

**Figure 13.**
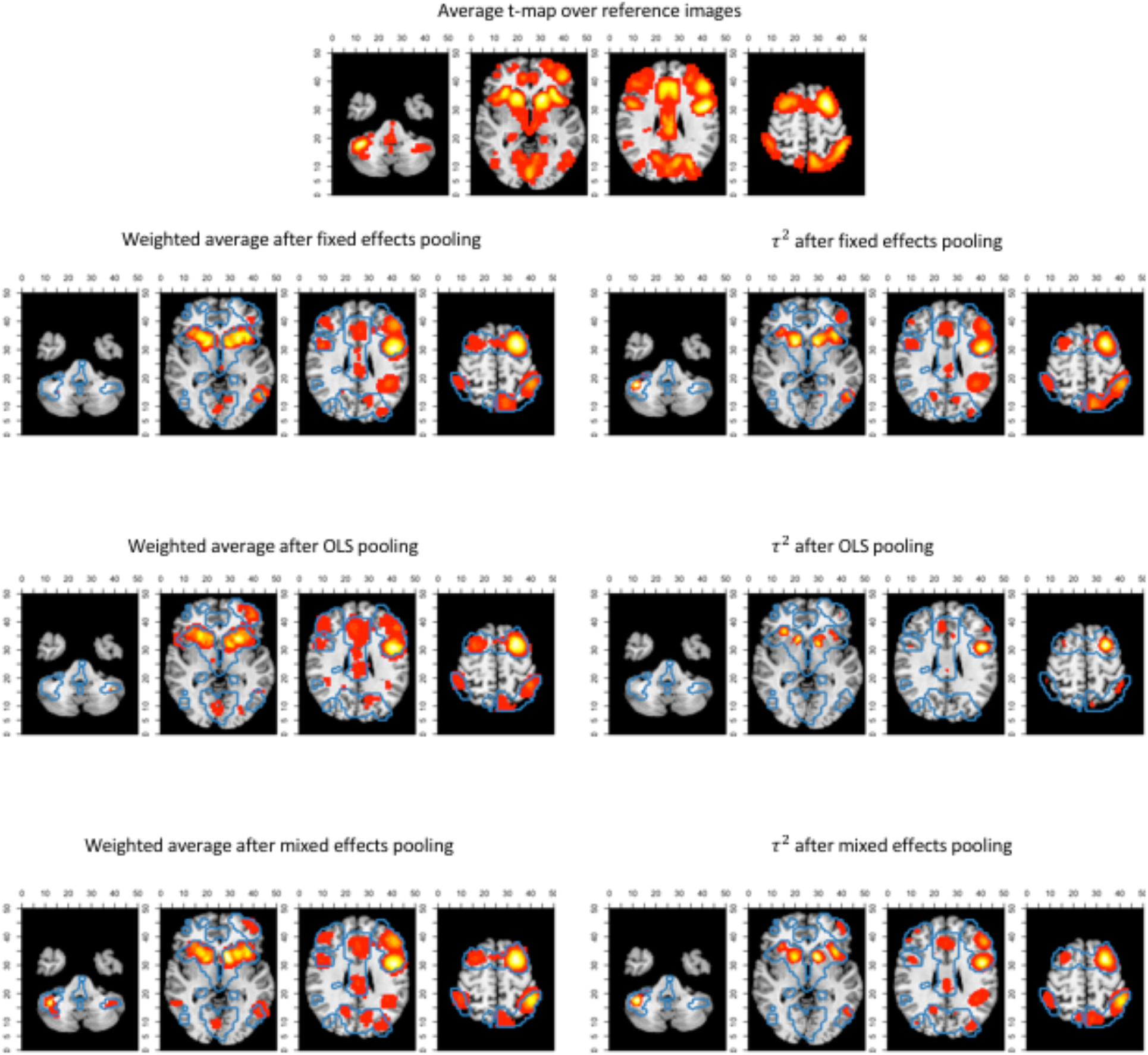
Slices (MNI z-coordinates from left to right: −44, −4,26 and 58) showing the average *t*-map of the reference images, the estimated variance between studies and the weighted average of the random effects meta-analysis (statistically significant voxels only) using the 3 pooling models for *K* = 10. The contour lines represent the average *t*-map of the reference images shown as illustration.

We observe the higher levels of between study heterogeneity mostly in the same regions that are statistically significant in the random (and fixed) effects meta-analysis (figure 13). OLS pooling generates less between study heterogeneity compared to fixed and mixed effects pooling. This corresponds to the overall smaller differences in performance between fixed and random effects meta-analysis we observe when using OLS pooling (e.g. see figures 3 and 9).

## 4 Discussion

In this paper, we studied how (1) the balance between false and true positives and (2) activation reliability for various coordinate-based meta-analysis (CBMA) methods in fMRI is influenced by an analytic choice at the study level. We applied a resampling scheme on a large existing dataset (*N* = 1400) to create a test condition and an independent evaluation condition. Each test condition corresponds to a combination of (a) a method for pooling subjects within studies and (b) a meta-analytic method for pooling studies. For (a), we considered ordinary least squares, fixed effects and mixed effects modelling in FSL and for (b) we considered an activation likelihood estimation (ALE), a fixed effects coordinate-based meta-analysis and a random effects coordinate-based meta-analysis. We generated meta-analyses consisting of either 10, 20 or 35 studies. The evaluation condition corresponded to a high-powered image that was used as a reference outcome for comparison with the meta-analytical results.

Comparing the test and evaluation condition enabled to calculate false and true positive hits of the metaanalyses depicted in ROC curves for each specific combination. By resampling within test conditions, we explored various measures of reliability.

In our study, we found the most optimal balance between false and true positives when combining a mixed effects group level model with a random effects meta-analysis. For less than 20 studies in the meta-analyses, adding more studies lead to a better balance for this analysis pipeline. When the meta-analysis contained at least 20 studies, there was no further considerable improvement by adding studies. Our results further indicate that the combination of a random effects meta-analysis performed better with respect to activation reliability when combined with a fixed or mixed effects group level model. There are however two disadvantages when using fixed effects group level models. First, inference is restricted to the participants included in the study (Mumford and Nichols, 2006). Second, it has been shown that fixed effects models tend to be liberal (Mumford and Nichols, 2006). Hence, comparing two images with a large amount of positive hits (either be true or false positives) likely corresponds with an increased overlap.

Noticeably, the ROC curves demonstrate a worse balance between false and true positives when OLS group level models are used to pool subjects within studies, regardless of the meta-analysis. As shown in Mumford and Nichols (2009), OLS models tend to be associated with conservative hypothesis testing and a loss of power depending on the sample size and the extent to which the assumption of homogeneous within subject variability is violated (see also Friston et al. 2005). Our results are in line with Roels et al. (2016) who show favourable ROC curves in parametric testing of the mixed effects group level model compared to OLS.

Regarding CBMA, it can be noted that even though ALE only includes peak location and not peak height (effect size), results converge to the same brain regions associated with high effect sizes in the reference images. Subsequently, the ALE results tend to involve brain regions that correspond to the detected regions in the reference images. Our observations are in line with Eickhoff et al. (2016b) in the sense that ALE meta-analyses require at least 20 studies. At this point, the outcome with respect to the ROC curves are close to the fixed and random effects methods for CBMA. These findings differ from Radua et al. (2012), who observe much lower values for sensitivity when comparing ALE to seed based *d*-mapping. Their study was limited however to 10 studies per meta-analysis. Furthermore, these authors applied a false discovery rate correction in ALE (at level 0.05) which is shown to be relatively low in sensitivity and susceptible to spurious activation for ALE maps (Eickhoff et al., 2016b). We on the other hand looked at a range of false positive rates given a significance level *α* which enables to study the power of procedures at an observed false positive rate.

We observed a lower reliability when using ALE compared with the fixed and random effects methods for CBMA, even when 35 studies were included in the meta-analysis. We propose the following explanations. First in low study set sizes and as shown in Eickhoff et al. (2016b), ALE results that include only 10 studies are more likely to be driven by one single experiment. Second, the two approaches differ in the kernel sizes when modelling the foci. As described in Radua et al. (2012) and Eickhoff et al. (2009), the ALE algorithm relies on kernels with a smaller full-width at half maximum than the fixed and random effects meta-analyses. This results in a greater number of small clusters of activation when using ALE. These images are more prone to be a hit or miss in a replication setting, depending on the sample size and the observed effect size. Third, the various methods use different approaches to correct for the multiple testing problem. For ALE we used the cFWE correction that was extensively validated in Eickhoff et al. (2016b). The fixed and random effects CBMA was implemented using the recommended thresholding of seed based d-mapping that relies on two (uncorrected) thresholds rather than explicitly correcting *P*-values. It remains unclear how this two-step thresholding procedure behaves in a range of scenarios where both the amount and location of peaks with respect to the true effect varies strongly.

We conclude with discussing some shortcomings of this paper.

First, we did not investigate adaptive smoothing kernels such as the anisotropic kernel described in Radua et al. (2014). This type of kernel incorporates spatial information of the brain structure. These kernels are promising as they potentially result in a better delineation of the activated brain regions in a meta-analysis rather than the Gaussian spheres we observed in our results.

Second, our results are characterized by low between-study heterogeneity since each study is created by sampling from the same dataset. In a real meta-analysis, we expect higher between study variability as it will include studies with a range of different scanner settings, paradigm operationalisations and sample populations. In previous versions of this manuscript, we tested (1) sampling subjects in figure 2 according to the scanning site involved in the IMAGEN project and (2) clustering subjects based on their individual effect size maps into individual studies to achieve higher between-study variability. However, these design adaptations did not yield substantial higher between-study heterogeneity.

Third, we limited our comparison to a fixed and random effects model implementation of an effect size based CBMA method with ALE, the most used CBMA method that only uses peak location. There are alternatives for ALE that also only use the location of local maxima such as Multilevel Kernel Density Analysis (Wager et al, 2007, 2009).

Fourth, we did not explicitly investigate the influence of the sample size of individual studies on the outcome of a meta-analysis. However, Tahmasebi et al. (2012) used the same IMAGEN dataset (though with a different contrast) to measure the effect of the sample size on the variability of the locations of peak activity in group analyses (study level). Their results indicate that 30 participants or more are needed so that locations of peak activity stabilize around a reference point. For similar results, see Thirion et al. (2007) who recommend at least 20 participants in a group analysis to achieve acceptable classification agreement. This was defined as the concordance between group analyses containing different subjects performing the same experimental design on declaring which voxels are truly active.

Finally, it should be stressed that our study does not reveal which combinations are more robust against the presence of bias. This bias could include (1) publication bias (Rothstein et al., 2005), (2) bias due to missing information since only statistically significant peak coordinates and/or peak effect sizes are used within studies and not the entire image, (3) or in the case of effect size based CBMA bias due to missing data if peak effect sizes for some studies are not reported (Costafreda, 2009; Wager et al., 2007). Seed based d-mapping, uses imputations to solve this latter missing data problem. As we did not have any missing data in our simulations, we did not evaluate the influence of these missing data on the performance of the various CBMA methods.

## 5 Conclusion

There is a clear loss of information when fMRI meta-analyses are restricted to coordinates of peak activation. However, if complete statistical parametric maps are unavailable, then coordinate based meta-analyses provide a way to aggregate results. We have investigated the trajectory of fMRI results from the choice of statistical group model at the study level to different coordinate-based meta-analysis methods. Our results favour the combination of mixed effects models in the second stage of the GLM procedure combined with random effects meta-analyses which rely on both the coordinates and effect sizes of the local maxima. Our results indicated (1) a higher balance between the false and true positive rate when compared to a high-powered reference image and (2) a higher reliability if the meta-analysis contains at least 20 or 35 studies. The popular Activation Likelihood Estimation method for coordinate-based meta-analysis provides a slightly lower but still comparable balance between false and true positives. However, it needs at least 35 studies to approach the higher levels of reliability associated with a random effects model for coordinate-based meta-analysis. The main advantage of our work consists of using a large database, while the main limitation is the restriction to only one dataset. We argue that this work provides substantial insight into the performance of coordinate based meta-analyses for fMRI.

## Acknowledgments

We would like to thank Jean-Baptiste Poline for the many, fruitful comments and discussions on this study. The computational resources (Stevin Supercomputer Infrastructure) and services used in this work were provided by the VSC (Flemish Supercomputer Center), funded by Ghent University, the Hercules Foundation, and the Flemish Government department EWI. Ruth Seurinck and Beatrijs Moerkerke would like to acknowledge the Research Foundation Flanders (FWO) for financial support (Grant G.0149.14).

Furthermore, this work received support from the following sources: the European Union-funded FP6 Integrated Project IMAGEN (Reinforcement-related behaviour in normal brain function and psychopathology) (LSHM-CT-2007-037286), the Horizon 2020 funded ERC Advanced Grant ‘STRATIFY’ (Brain network based stratification of reinforcement-related disorders) (695313), ERANID (Understanding the Interplay between Cultural, Biological and Subjective Factors in Drug Use Pathways) (PR-ST-0416-10004), BRIDGET (JPND: BRain Imaging, cognition Dementia and next generation GEnomics) (MR/N027558/1), the FP7 projects IMAGEMEND(602450; IMAging GEnetics for MENtal Disorders) and MATRICS (603016), the Innovative Medicine Initiative Project EU-AIMS (115300-2), the Medical Research Council Grant ‘c-VEDA’ (Consortium on Vulnerability to Externalizing Disorders and Addictions) (MR/N000390/1), the Swedish Research Council FORMAS, the Medical Research Council, the National Institute for Health Research (NIHR) Biomedical Research Centre at South London and Maudsley NHS Foundation Trust and King’s College London, the Bundesministeriumfür Bildung und Forschung (BMBF grants 01GS08152; 01EV0711; eMED SysAlc01ZX1311A; Forschungsnetz AERIAL), the Deutsche Forschungsgemeinschaft (DFG grants SM 80/7-1, SM 80/7-2, SFB 940/1). Further support was provided by grants from: ANR (project AF12-NEUR0008-01 - WM2NA, and ANR-12-SAMA-0004), the Fondation de France, the Fondation pour la Recherche Médicale, the Mission Interministérielle de Lutte-contre-les-Drogues-et-les-Conduites-Addictives (MILDECA), the Assistance-Publique-Hôpitaux-de-Paris and INSERM (interface grant), Paris Sud University IDEX 2012; the National Institutes of Health, U.S.A. (Axon, Testosterone and Mental Health during Adolescence; RO1 MH085772-01A1), and by NIH Consortium grant U54 EB020403, supported by a cross-NIH alliance that funds Big Data to Knowledge Centres of Excellence.

## Author Contributions

H.B., R.S., S.K. and B.M. contributed to the conception and design of the manuscript. Data collection and single subject analyses were carried out by the IMAGEN consortium represented by T.B., G.B., A.L.W., B.D., J-L. M., H.L., T.P. and S.M.D. Data analysis and interpretation for this study was performed by H.B., R.S. and B.M. Next, H.B. developed the initial draft of the manuscript. Finally, all authors approve the version to be published.

## Appendix

### 1. Distributions of amount and cluster sizes

For *K* = 10, 20 and 35, we plot the amount of overlapping and unique clusters with the cluster sizes (expressed in number of voxels) next to it. This is calculated on the pairwise comparisons of the *I* unique iterations. We plot the results for each group level model and CBMA.

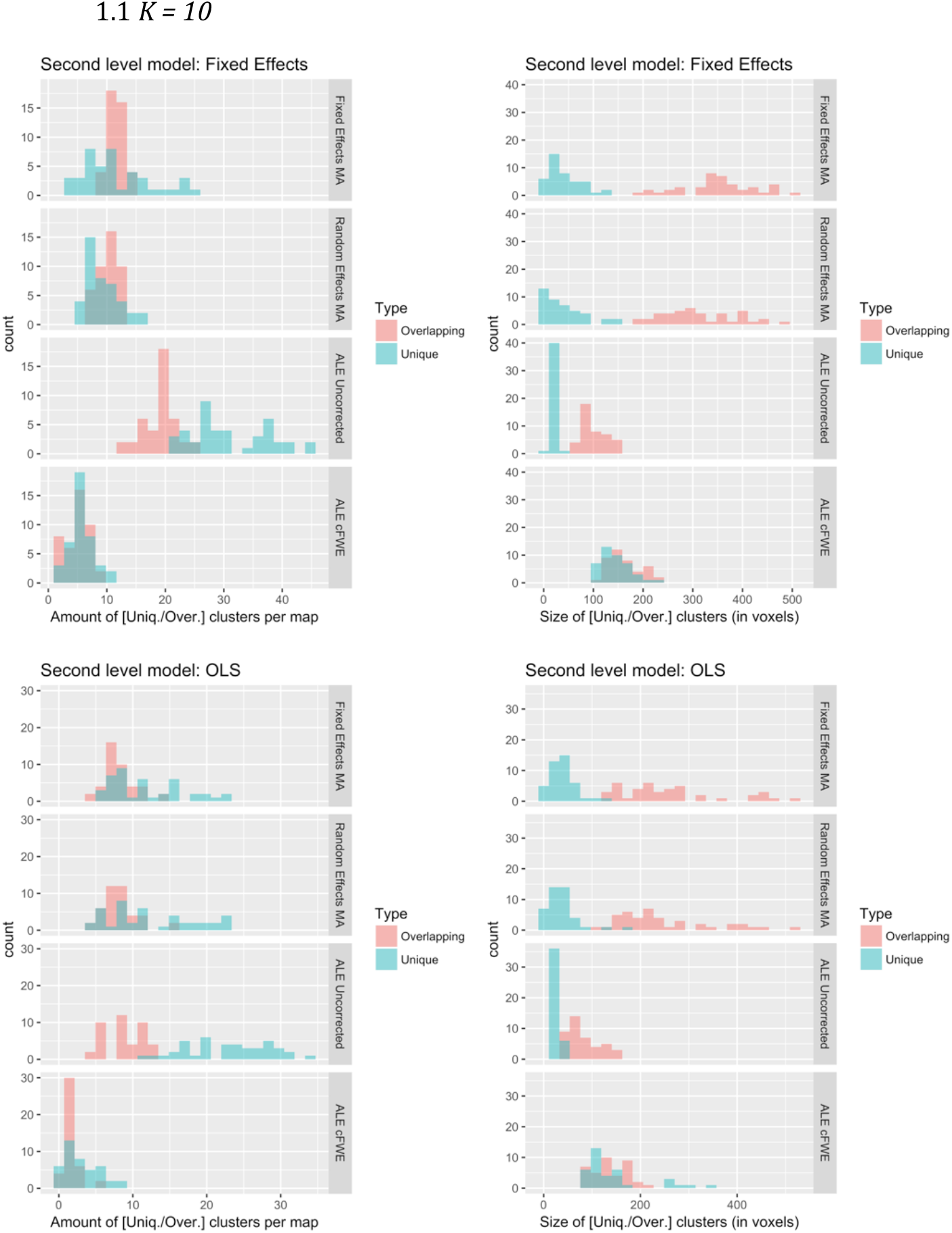
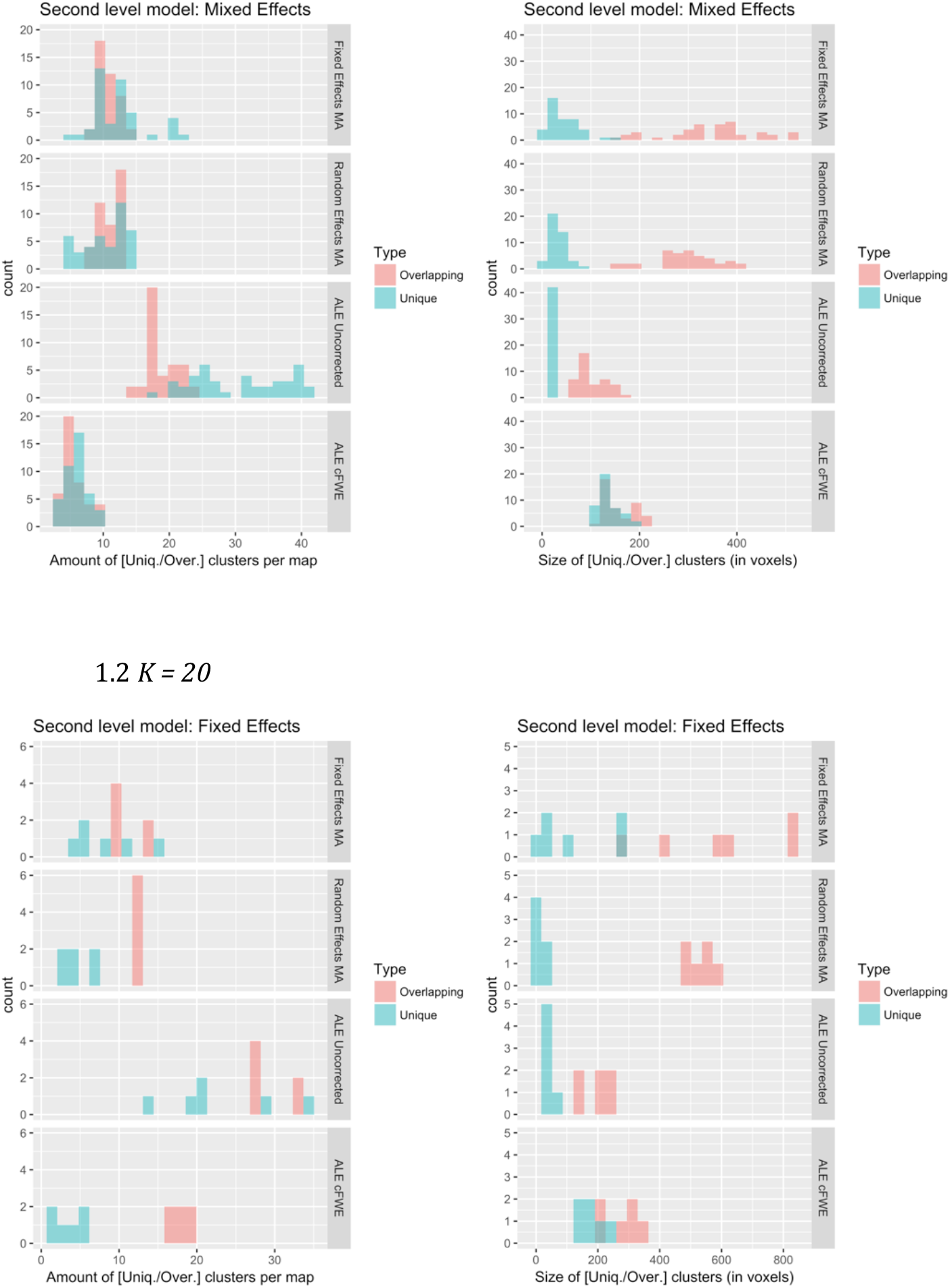
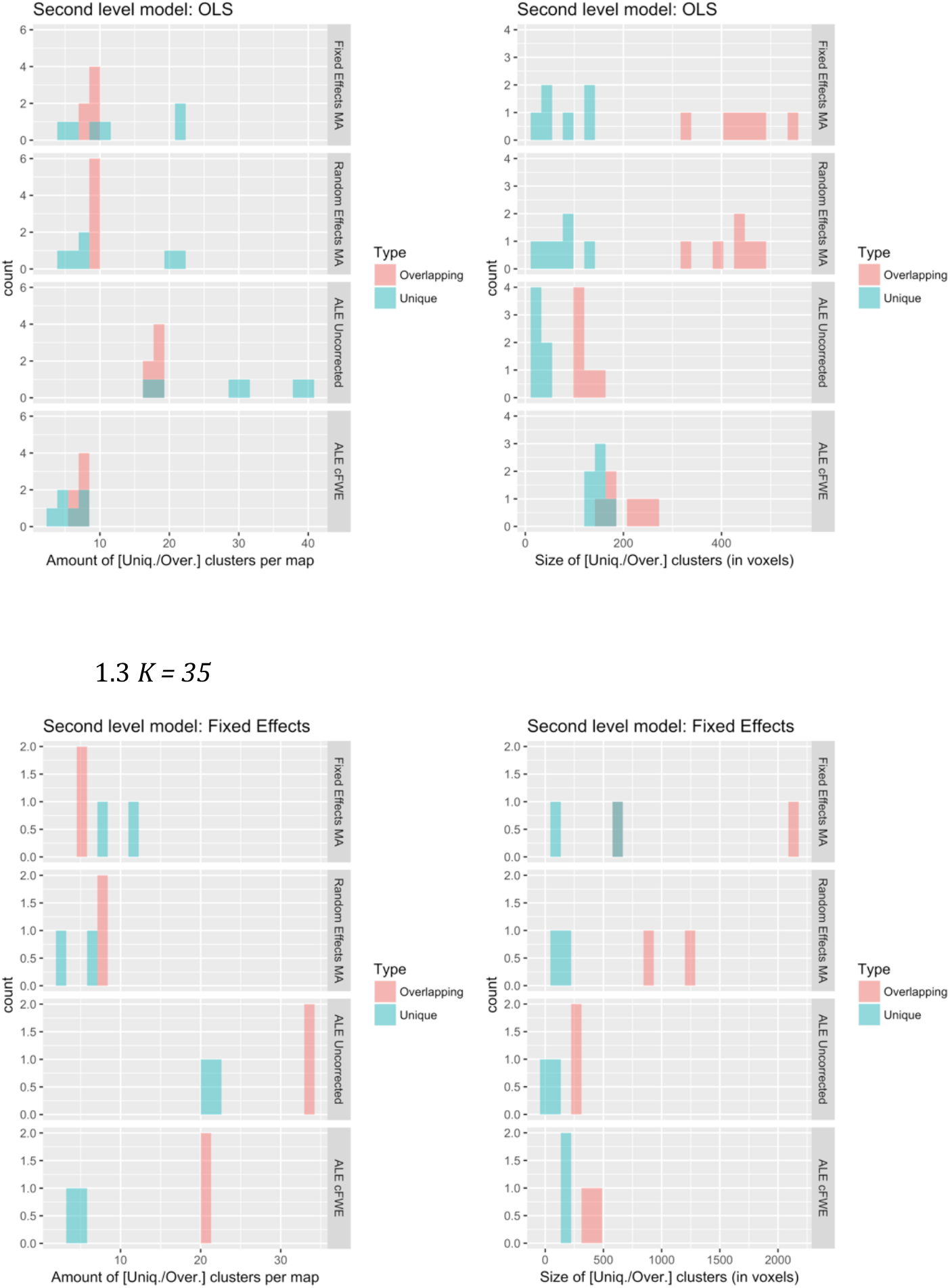
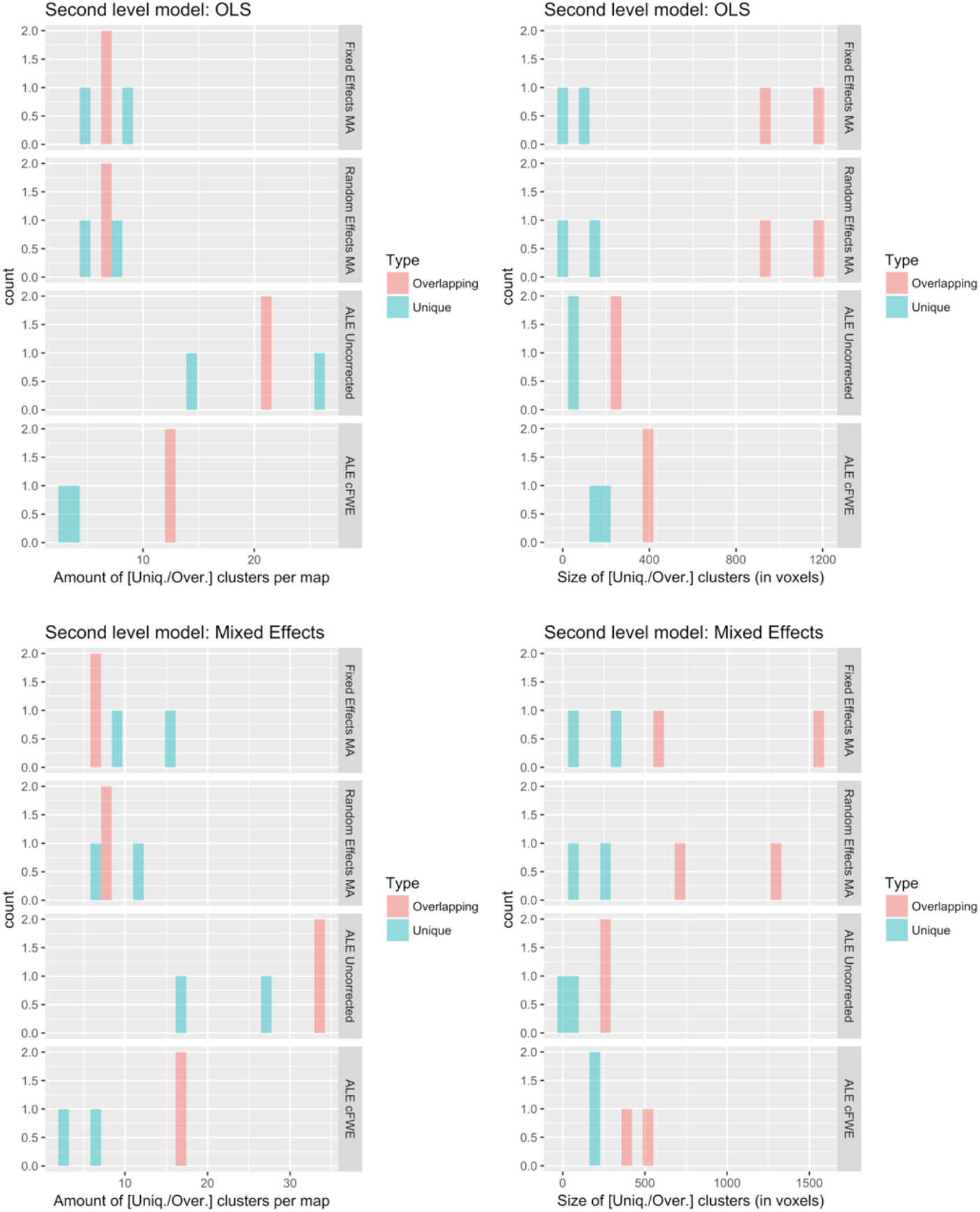

